# A universal system for streamlined genome integrations with CRISPR-associated transposases

**DOI:** 10.1101/2022.05.30.494051

**Authors:** Megan Wang, Charles Sanfiorenzo, Raymond J. Zhang, Kaihang Wang

## Abstract

Genome engineering tools in bacteria are limited by their targeting abilities, cargo size capacities, and integration efficiencies. Programmable Cas-directed transposons have been shown to bypass these constraints; however, genome integrations with these Cas-directed transposons require a cargo plasmid carrying the desired DNA payload flanked by directed repeat transposon arms. This cloning pre-requisite significantly hinders the modularity and streamlining capabilities of Cas-directed transposon systems, diminishing their utility for genome engineering. Here, we present a system that can robustly integrate a linear DNA payload into the genome of *E. coli* by employing a Type I-F CRISPR-associated transposon from *Vibrio cholerae*. This system bypasses the traditional limiting factors of Cas-directed transposons by leveraging oligonucleotide design and nested polymerase chain reactions to reconstitute the transposon arms onto a designated cargo. Our findings demonstrate that this programmable linear integration method has high efficiencies in integrating large DNA payloads across multiple genomic loci.

## Introduction

Genome engineering technologies have been employed to program bacteria for a variety of applications. The development of these tools has allowed for the expression of heterologous genes in bacteria for the commercial production of industrial and medically relevant compounds^1*-2,3*,^ as well as for the discovery of novel metabolites ^4,5^. Oftentimes, genome integration is preferred over plasmid-based expression as the former bypasses the need for sustained antibiotic selection, enables tight control of gene expression levels through a stable chromosomal copy number, and lowers the metabolic burden on the cell. While integrating gene inserts into bacterial genomes is widely utilized and especially important in the field of synthetic biology, modifications to the genome on the kilobase scale remain difficult, typically requiring careful cloning and validation of donor DNA. Preeminent methods for genome integrations in *E. coli* include the use of homology-directed recombination^6,7^, site-specific recombinases^8,9^, and pseudo-random transposases^10,11^.

State of the art for homology-directed recombination, or “recombineering”, features the insertion of DNA mediated by ≥50-bp homology arms with the aid of bacteriophage recombinase proteins^12^. The most commonly used recombineering protein system in *Escherichia coli* (*E. coli*) is the Lambda (λ) - Red system derived from bacteriophage λ. The Lambda-Red recombineering system consists of three proteins: alpha, beta, and gamma^13^. Alpha, commonly known as the exonuclease (exo), resects the 5’-ended strand of dsDNA to expose a single-stranded region. This allows beta to bind to the single-stranded overhangs and guide the exposed end of an incoming piece of newly generated ssDNA to a homologous locus and facilitate homologous recombination. Gamma functions as an accessory protein, inhibiting endogenous RecBCD and SbcCD exonuclease activity to prevent digestion of the exposed recombination template. When expressed in *E. coli* cells, the Lambda-Red recombineering system permits the use of short regions of homology (∼50-100bp) to facilitate recombination between a target insert and a genomic locus^12,14^. Such homologous overhangs can easily be included within synthetic oligonucleotides used during DNA amplification through Polymerase Chain Reaction (PCR)^6^. This feature allows for the modular design of inserts targeted to specific genomic loci and eliminates the need for additional cloning steps characteristic of genome integration methods mediated by site-specific recombinases^15,16^. However, recombineering systems face a stringent upper size limit for the length of DNA that can be feasibly inserted into the genome of bacteria following intake of recombinant DNA.

In practice, recombination frequencies sharply decline when inserting cargos greater than 3-4kb^17^. To circumvent this pitfall, the Lambda-Red recombineering system has been coupled with CRISPR-Cas9 to expose target loci for recombineering through Cas9-induced double-stranded breaks in four different positions – 2 cuts in the genome, and 2 cuts in an artificial bacterial chromosome (BAC) carrying a synthetic DNA payload^17–20^. This concept, denoted in the literature as “replicon excision for enhanced genome engineering through programmed recombination” (REXER), enabled the serial integration and exchange of synthetic genomic segments onto the *E. coli* genome in 100-kb steps^17,19^. The use of REXER subsequently allowed for large chromosomal rearrangements and exchanges and ultimately resulted in the whole-genome synthesis of a synthetic 3.9-Mb Escherichia coli genome^18^. REXER, however, requires laborious cloning of vectors for both the synthetic BAC cargo and the Cas9 guide RNAs. Additionally, the lack of universal single-copy BAC backbones and recombineering proteins for bacteria^21–24^ are barriers to implementing REXER in many bacterial species.

Unlike canonical recombineering, the use of site-specific recombinases for genome engineering has been demonstrated to be agnostic to expression hosts and the lengths of DNA products utilized for genomic integration. Currently, serine and tyrosine recombinases, which are characterized by their catalytic site residues, are widely employed for genome engineering in bacteria^8,25,26^. Serine recombinases such as Bxb1^27^ and φC31^25,28^ integrases, attach to attB and attP sites to generate two double-stranded breaks creating 5′-phosphoserine bonds. Recombination of the two double-stranded break sites is directed by the formation of attL and attR sites. In contrast, tyrosine recombinases such as the Cre recombinase derived from the P1 phage, cleave single-strands to form 3’-phosphotyrosine bonds and are re-ligated through a Holliday junction-like intermediate^29,30^. Both tyrosine and serine recombinases can mediate deletion, inversion, and translocation events within single or multiple DNA molecules. While these features make site-specific recombinases an incredibly attractive tool suite for genome engineering operations, their use requires the existence of pre-engineered recombinase recognition sites at target genomic loci. This prerequisite greatly limits the modularity and streamlining capabilities of site-specific recombinases as genome engineering tools.

Transposases, such as Tn5 and Tn7, have also been used to integrate DNA into the genome^31–34^. Tn5 integrates nonspecifically in a “cut and paste” mechanism flanked by IS50R and IS50L, which are target sites of the Tn5 transposase^35^. The pseudo-random integration feature has been leveraged in a variety of applications, ranging from next-generation sequencing^36^ to gene essentially screens through random insertional mutagenesis^37^. In contrast, Tn7 transposes cargo at high frequencies downstream of the attTn7 site naturally located within many bacterial genomes^34^. Although Tn7 is useful for the high-efficiency integration of DNA payloads at a specific site, the inability to program insertion locations limits its genome editing capacities.

More recently, phylogenetic analyses have identified programmable Tn7-like elements that are evolutionarily linked to CRISPR-Cas systems^*38*^, now commonly referred to as CRISPR-associated transposons (CASTs). These CRISPR-associated transposase systems have been shown to deliver large DNA payloads and insert payloads ∼40-50-bp downstream of target sites at high efficiencies in *E. coli* in a CRISPR-RNA (crRNA) dependent manner^*39,40*^. The potential genome engineering capacities of two Tn7-like systems have been demonstrated. Isolated from *Scytonema hofmannii*, the ShCAST system is comprised of transposition proteins, TnsB, TnsC, TniQ, along with a Type V-K Cas12k DNA targeting protein. An orthogonal system discovered in *Vibrio cholerae* (*V. cholerae*), utilizes orthologous transposition proteins along with TnsA coupled to a Type I-F CRISPR-Cas Cascade (Cas5-8, Cas6, and Cas7) effector complex. Because the Type V-K Cas12k DNA targeting complex is devoid of a functional TnsA protein, which normally acts as a 5’ cleaving endonuclease, transposition from a plasmid donor onto the genome typically results in the integration of co-integrate products that include the donor plasmid backbone^*39,41–45*^. In light of this, the *V. cholerae* Type I-F CAST has been further utilized as a genome engineering tool for bacterial genomes. The INTEGRATE system, using the *V. cholerae* CAST, was designed for efficient on-target integration of large DNA payloads (up to 10-kb) within a single step^*46*^. The capstone of the INTEGRATE system is the all-in-one design containing the CAST, the donor DNA and the targeting crRNA on a single plasmid (pSPIN). While this system lays the groundwork for transitioning CASTs into robust genome engineering tools, state-of-the-art technologies for genome engineering using single plasmid INTEGRATE lack modularity and can be streamlined.

To address modularity and to further streamline capabilities of CASTs for bacterial genome engineering, we herein develop a system capable of transforming any DNA template into a compatible transposition insert for the *V. cholerae* Type I-F CRISPR-associated transposon. Previous work has included the Right (R) and Left (L) arms of the Tn7-like transposon on cargo vectors, requiring an intermediate step where the insert is cloned into the plasmid backbone prior to integration^*46*^. Our innovation eliminates the need for molecular cloning through the introduction of the R and L arms as overhangs for linear amplicons for genome integration. Furthermore, the inclusion of the crRNA guide onto a modular, easy-to-remove vector, allows rapid interchangeability of the target integration site as new cargo is introduced into the cell. With our innovations, this CAST system has the prospect to be as versatile as Lambda-Red recombination, but importantly, includes the ability to integrate large (∼10-kb) DNA payloads at efficiencies comparable to small (≤3-kb), linear cargo. Although not explored in this work, we expect the herein described amplicon-based CAST integration approach to be extendable towards more recently uncovered Type I-F CRISPR-associated transposons^*47*^, further simplifying the screening process for the functionality of orthogonal CASTs.

## Results and Discussion

### Design of Universal CAST Inserts via Nested Amplification and Inducible CRISPR-associated Transposition

The addition of the transposon’s R and L arms onto a linear DNA payload requires two sets of primer oligos including annealing/priming regions and segments of the transposon arms (Figure 1a-b). We decided to reconstitute a full-length L arm (145-bp) and a truncated R arm (57-bp vs 125-bp) through the primer overhangs based on previous work^40^, where it was documented that the use of a truncated R arm led to higher transposition efficiencies and additionally increased the RL::LR orientation bias observed at the integration site. Given that the entirety of the L arm exceeds the maximum oligonucleotide length offered by most commercial suppliers, two rounds of amplification were necessary to successfully reconstitute the L arm in the cargo amplicon. An added advantage of this nested PCR method is the extreme dilution of the cargo’s plasmid template, resulting in no detectable plasmid-harboring colony-forming units (CFUs) following transformation^48^. In order to select for genomic insertion, a cargo encoding for at least one positive marker (+1) was utilized as the template for nested amplification (Figure 1a).

**Figure 1.**
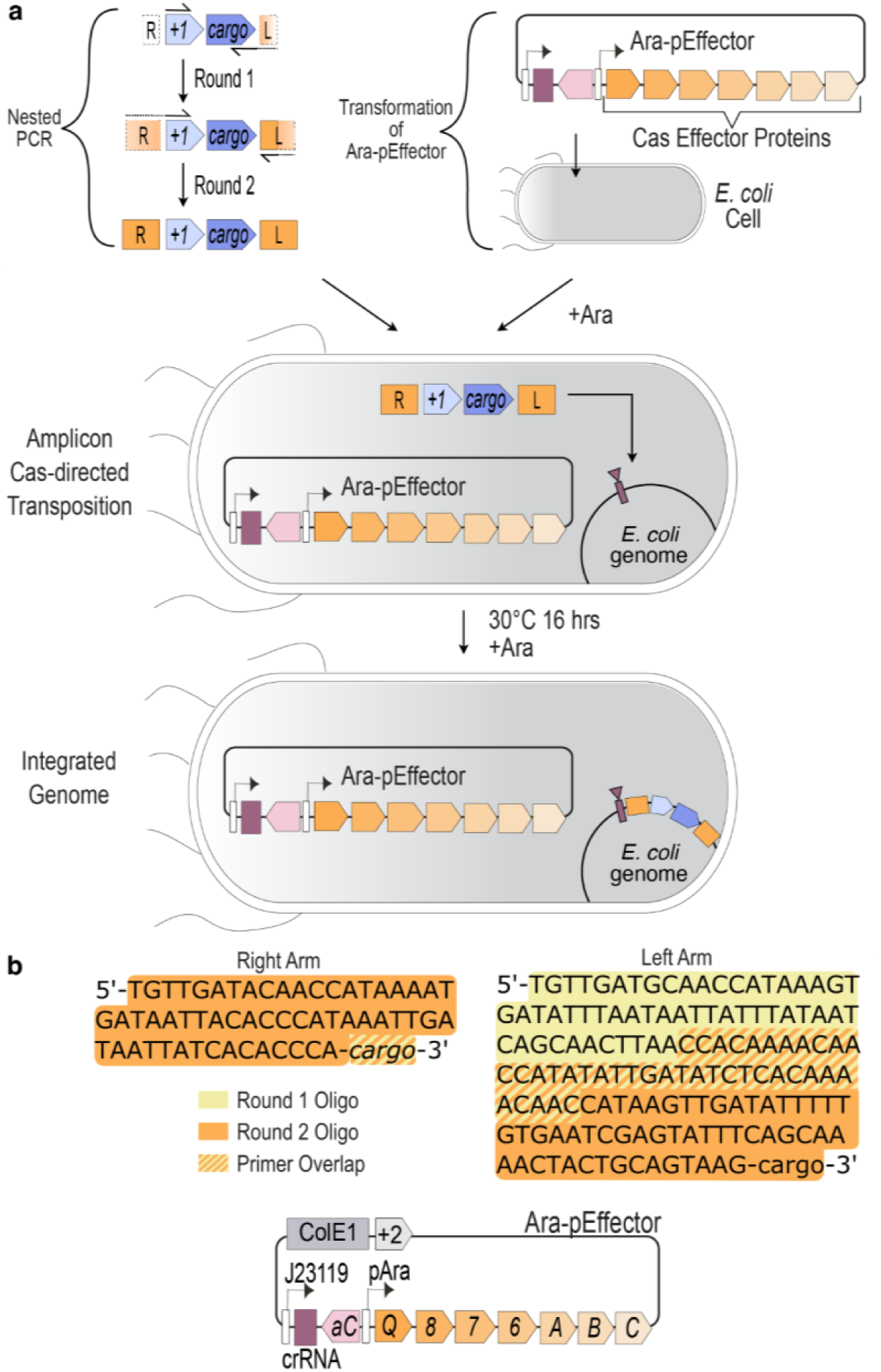
Streamlined genomic insertion through nested amplification and transformation of transposable cargo. (A) Nested PCR method for reconstituting transposon arms onto the cargo and transformation of the Ara-pEffector for cargo integration in the genome. Cells are pre-induced with arabinose prior to the transformation of the final transposable amplicon. Following transformation, cells are recovered and induced for a period of 16 hours at 30º C before plating in solid LB Agar media. A positive (+1) selection marker encoded within the cargo is used to select for genomic integration. (B) Detailed design of nested PCR amplification process of the insert cargo and schematic of Ara-pEffector construct. The first round of nested amplification has both primers annealing to the cargo template, with one primer containing 91-bp of the L arm. The second round has a primer overlapping the cargo region containing the full R arm and a primer overlapping the last 38-bp resulting from the first round’s L arm primer, completing the last 54-bp of the L arm through the primer’s overhang. The Ara-pEffector plasmid, carried under the ColE1 backbone with a positive selection marker (+2), consists of the VchCAST Effector proteins induced by the arabinose toggable pAra/pBAD promoter and a targeting crRNA.

To establish togglable control over the *V. cholerae* Type I-F CAST proteins used in this system, an arabinose-inducible and glucose-suppressible CAST construct, termed Ara-pEffector, was built off the constitutively-expressed VchINT pEffector construct described in the INTEGRATE system^46^ (Figure 1b). The pEffector plasmid is carried under the high-copy (∼15-20 plasmid copies) ColE1 backbone, with a guide crRNA expressed through a constitutive, synthetic J23119 promoter to ensure that sufficient crRNA is present in the cell for adequate recruitment and assembly with the CAST proteins in anticipation of transforming the final PCR product. To select for cells harboring the Ara-pEffector, the construct additionally encodes for a spectinomycin resistance gene (SmR). Once generated, the final linear amplicon with reconstituted L and R transposon arms can be transformed into arabinose-induced bacteria harboring the Ara-pEffector. After recovering and inducing transformed cells for a period of 16 hours at 30 ºC, the cells are spread in selective LB Agar plates and incubated overnight at 37 ºC. Obtained CFUs can then be quantified and assessed for precise genomic integration. By coupling nested amplification of linear cargo to CAST-based genomic integrations, the system described in this work streamlines the cargo assembly process for programmable genome insertion.

### Transposition Efficiencies are Comparable Across Genomic Loci

To test our system and interrogate integration efficiencies in a genomic context, we designed four integration locations spread across the *E. coli* genome (Figure 2a). The crRNA target sites for the four depicted integration locations – TS1, TS2, TS5, and TS4 – were chosen on the assumption that they would yield genetically viable integration events (i.e. no essential information disrupted) based on experimental validation carried out from previous work^17,18^. Each crRNA target site is between 700-kb to 1.8-Mb from the adjacent target sites. For the linear DNA payload, we chose to integrate a positive/negative double-selection cassette consisting of the chloramphenicol resistance marker (+1, *CmR*) and the sacB negative selection marker conferring sucrose susceptibility. Both positive and negative markers would aid in phenotyping for genomic integrations. The linear DNA payload additionally contained a GFP gene, serving as a visual marker for successful genomic integration. The entire length of the DNA payload was consequently around 3-kb.

**Figure 2.**
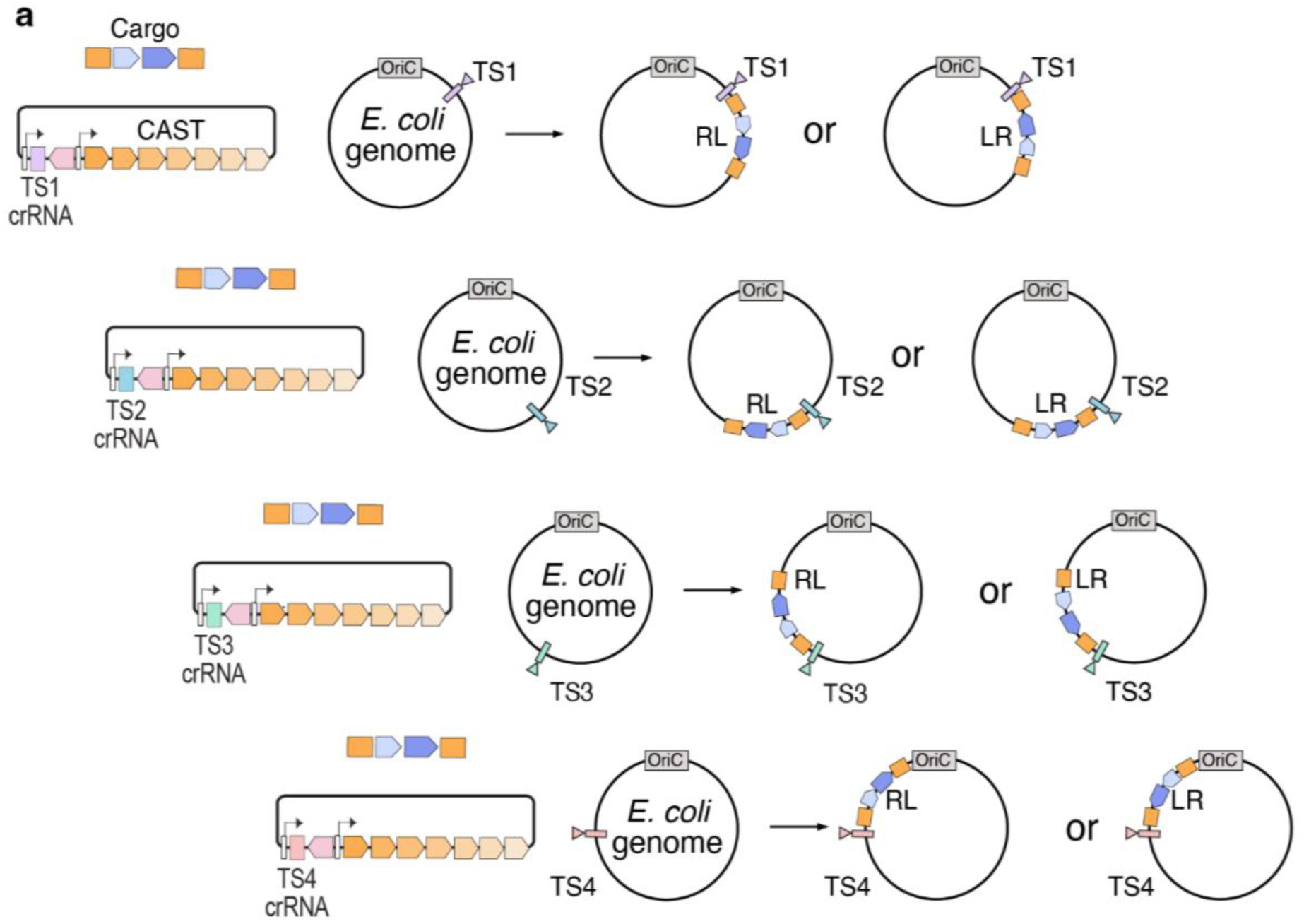
Targeting different genomic locations through interchanging crRNA guides on the Ara-pEffector plasmid. (A) Integration of cargo in both RL and LR orientations at the TS1, TS2, TS3, or TS4 site. Bacterial strains are generated carrying either Ara-pEffector^TS1^, Ara-pEffector^TS2^, Ara-pEffector^TS3^, or Ara-pEffector^TS4^ prior to transformation of amplified DNA payload product.

After transformation of the linear payload into *E. coli* DH10b (Invitrogen) cells containing a transiently induced Ara-pEffector with its respective crRNA, cells were recovered and plated onto selective agar plates, and GFP+ CFU were quantified. All CFUs detected in this assay exhibited GFP+ signal. These results showcase that integration efficiencies observed for the linear cargo were consistent across all four genomic loci (Figure 3a), suggesting the versatility of this PCR-based approach for programmable genome engineering. Although integration into TS1 and TS3 resulted in slightly lower efficiencies than integration into TS2 and TS4, the total amount of CFUs was comparable. Notably, the Ara-pEffectors with a non-targeting (NT) crRNA or with no crRNA yielded no colonies, demonstrating that integration of PCR products with reconstituted L-R transposon arms is dependent on crRNA-mediated targeting. Further downstream phenotypic screens were conducted via spotting assays on positive and negative antibiotic markers (Supporting Information, Figure S1a), and junctions resulting from integration events into each target were assessed via PCR-based genotyping (Supporting Information, Figure S1b). Both phenotypes and PCR genotypes were consistent for integration events for all clones screened in our assay. Sanger sequencing was performed on the genotyped replicates, further confirming successful integration into the respective target sites (Supporting Information, Figure S2).

**Figure 3.**
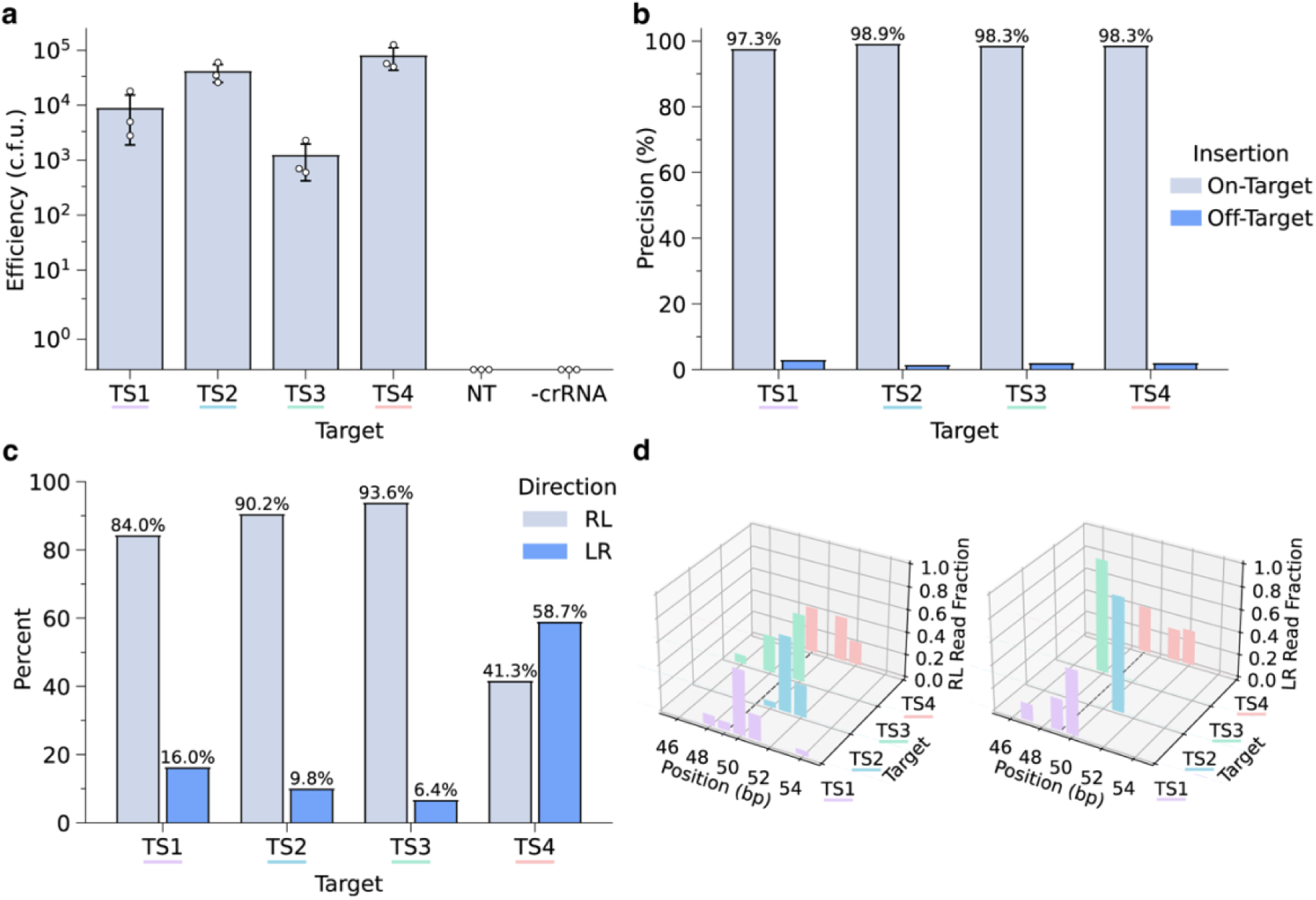
Analyses of integrations resulting from the nested amplification method targeted four genomic loci (TS1, TS2, TS3, TS4). (A) Integration efficiencies based on GFP+ CFU (biological replicates, n=3). (B) Integration accuracies, (C) directionality biases, and (D) insertion locations generated from amplicon sequencing of bacterial populations. The dashed line in (D) represents the most frequently integrated site (49-bp after the crRNA target) reported in previous studies. Bars indicate the mean of biological triplicates with the error bars representing the standard error.

To further assess integration efficiencies and to inspect bias based on integration distance from the target site and insertion orientation, amplicon sequencing was performed on populations resulting from integration experiments into TS1, TS2, TS3, and TS4 targeting sites (Figures 3b-d). Three sites (TS1, TS2, and TS3) follow the previously described insert directionality bias (>75% RL orientation)^*46*^; however, the TS4 integration site showed a slightly greater LR orientation bias (Figure 3b). Previous literature on bacterial transposon Tn7 has reported that the directionality bias of Tn7 is dependent on insertion location relative to the OriC and Ter sites^*49*^. These findings elucidate the possibility of there being additional directionality biases in the genomic context for integrations mediated by the *V. cholerae* Type I-F CAST proteins. Further experimentation is required to properly interrogate this claim. On-target site-specificity is defined as an insertion between −10-bp and +10-bp from the site 50-bp downstream of the crRNA target sequence. Using this metric, on-target insertion accuracy was assessed to be >97% for all four RNA crRNA guide targets (Figure 3c). The exact insertion location sorted by the directionality of the DNA payload was also computed, wherein most integrations occurred between 48-bp and 51-bp downstream from the target site (Figure 3d), as described in previous literature^,*40,46*^. Read coverage across insertion sites was plotted (Supporting Information, Figure S3).

### CAST integrations outcompete Lambda-Red mediated genomic integrations in respect to target site, insert length, and sequence identity

Following characterization of integration efficiencies, accuracies, and biases for a 3-kb transposon insert across four genomic loci, we wanted to directly compare its efficiencies to conventional Lambda-Red recombineering (Figure 4a). Both systems were tested to integrate three payloads of various sizes: 1-kb, 3-kb, and 10-kb, each containing a selection cassette driven by an EM7 promoter and conferring chloramphenicol resistance. The largest, 10-kb insert included an additional bacterial chemiluminescence (luxABCDE) cassette. Reconstitution of the R and L transposon arms onto the linear amplicon was completed through the nested PCR method described in this work.

**Figure 4.**
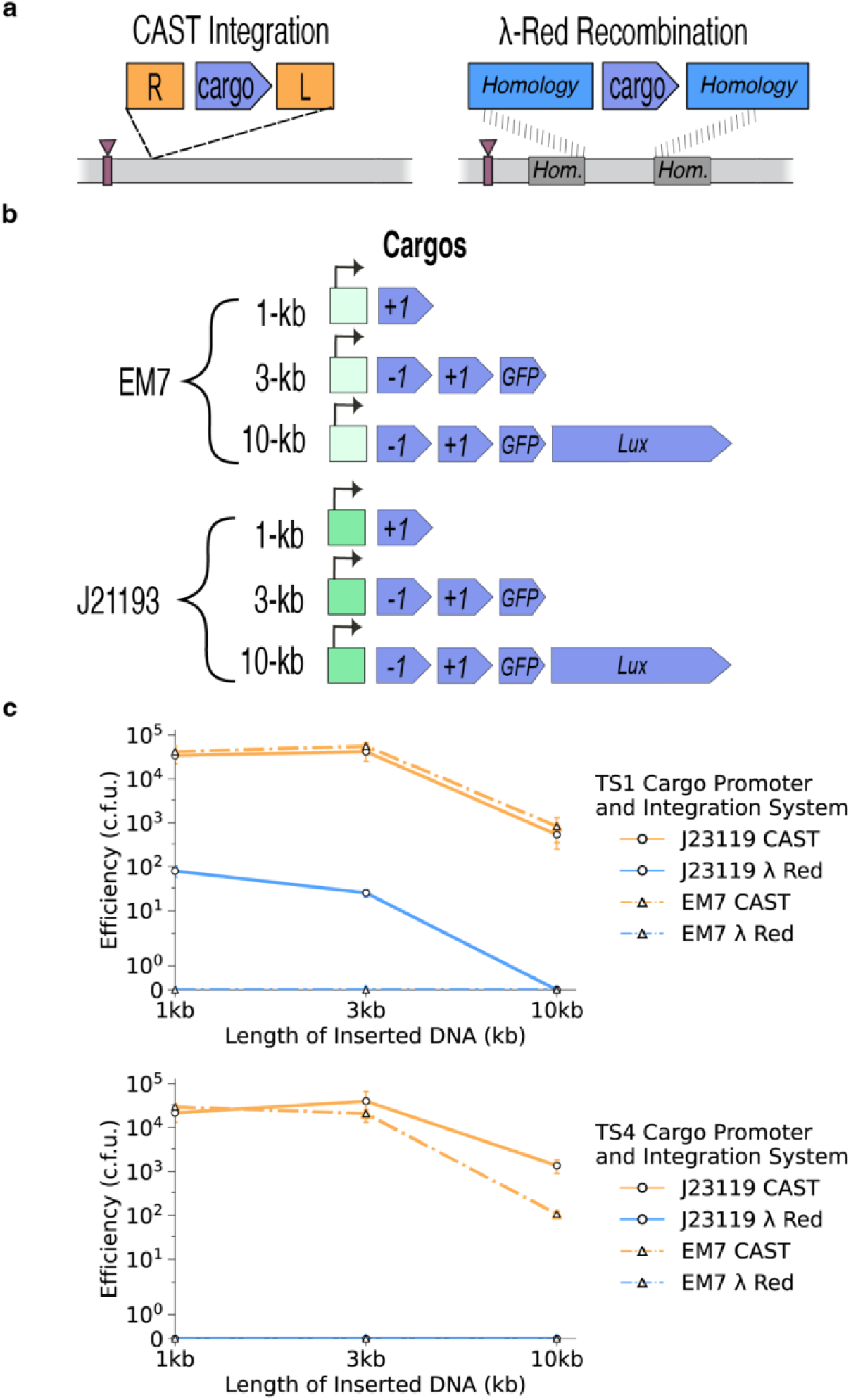
Comparison of linear CAST-directed and Lambda-Red mediated integration efficiencies. (A) Schematics of CAST-directed integration and Lambda-Red recombination of cargo. (B) Linear cargos of sizes 1-kb, 3-kb, and 10-kb driven by the EM7 or J23119 promoter were integrated into two targets. (C) TS1 integration via linear CAST was between 2 to 3 orders of magnitude more efficient than Lambda-Red recombination for 1-kb and 3-kb inserts. The 10-kb insert did not yield colonies for both target sites tested. TS4 integrations through Lambda-Red failed for all cargo sizes tested, while CAST successfully integrated identical cargos into TS4. Bars indicate the mean of biological triplicates with the error bars representing the standard error.

Since the Lambda-Red system has the ability to precisely target integration down to the single base-pair resolution, the three payloads were integrated into a site 50-bp downstream of the corresponding TS1 and TS4 crRNA targets on the genome. The homology regions included in the oligos for Lambda-Red recombineering were designated as the 50-bp directly upstream and downstream of the expected CAST integration site. Lambda-Red-mediated recombination failed to integrate all three EM7-driven payloads (1-kb, 3-kb and 10-kb) used for earlier CAST-driven integrations (Figure 4c). These recombination experiments were repeated on multiple occasions, and by multiple authors in this work. These results were surprising since linear cargos the size of 1-3-kb is expected to fall within the working range for Lambda-Red-based recombinations. We reasoned that for Lambda-Red insertions, the specific nucleotide sequence of the linear donor DNA might decrease, or even nullify, genomic integration efficiencies.

Specifically, we hypothesized that the EM7 promoter sequence driving the positive/negative marker cassette was hindering integration efficiencies due to possessing partial homology to untargeted loci in the genome. To test this hypothesis, we replaced the EM7 promoter with a synthetic J23119 promoter and re-attempted Lambda-Red integrations (Figure 4b). Replacement of the EM7 promoter with the synthetic J23119 promoter circumvented difficulties in Lambda-Red-mediated integrations into some, but not all, target sites tested. Our results show that efficiencies for Lambda-Red recombination for the 1-kb and 3-kb cargos into the TS1 target site are around 2-3 fold lower than CAST-driven linear integration while the 10-kb cargo failed to integrate (Figure 4c). Lambda-Red recombination into the TS4 site failed for all three cargo sizes (Figure 4c). These results showcase that CAST-driven integrations can provide flexibility in respect to genomic integration locations, agnostic of sequences encoded within cargos.

In comparing integration efficiencies between the Lambda-Red and CAST-driven systems, integration of the 10-kb cargo with the CAST method showed comparable efficiencies to the 1-kb and 3-kb cargos. The luxABCDE luminescence cassette was used to screen for full-length 10-kb integrants. After transformation of the linear payload (either 1-kb, 3-kb, or 10-kb) into pre-induced *E. coli* DH10b cells containing the respective Ara-pEffector (TS1 or TS4), cells were plated following recovery onto selective agar plates. Chloramphenicol resistant, GFP+, and bioluminescent CFUs were subsequently quantified. For both genomic targets tested for the 10-kb insert, all CFUs observed in the selective agar plates exhibited both GFP and luminescent positive signals, indicating full-length integration. A ∼1.5 fold-change drop was observed when comparing integration efficiencies quantified for the 10-kb versus the similarly efficient 1-kb and 3-kb inserts, with the shorter inserts showcasing higher integration efficiencies (Figure 4c). The decrease in integration efficiency observed for the 10-kb insert is likely due to decreased transformation efficiencies as a function of PCR product length, as commonly observed in the electroporation procedure utilized in this work (See Materials and Methods). Nonetheless, despite this drop in integration efficiency for a longer payload, integration of the 10-kb insert yielded colonies on the 10^2^ to 10^3^ scale. These trends in integration efficiencies for the larger, 10-kb payload was consistent across EM7 and J23119-driven cargos.

A subset (n=8 per target locus) of CFUs utilized for quantification of integration efficiencies were resuspended and screened phenotypically and genotypically to further validate successful integration events (Figure 5a-b). Phenotypic screens were conducted via spotting assays with plates containing spectinomycin to select for cells containing the Ara-pEffector in combination with other selection drugs. Successful 10-kb integrants are expected to be resistant to chloramphenicol, susceptible to sucrose, and both GFP+ and bioluminescent, while successful 3-kb integrants should exhibit the same growth selection but should not luminesce. *E. coli* DH10b cells harboring the corresponding Ara-pEffector were spotted alongside integrants as phenotyping controls for resistances, susceptibilities, and fluorescence/luminescence. Cells only harboring the Ara-pEffector plasmid are expected to be GFP-, not bioluminescent, and susceptible to chloramphenicol.

**Figure 5.**
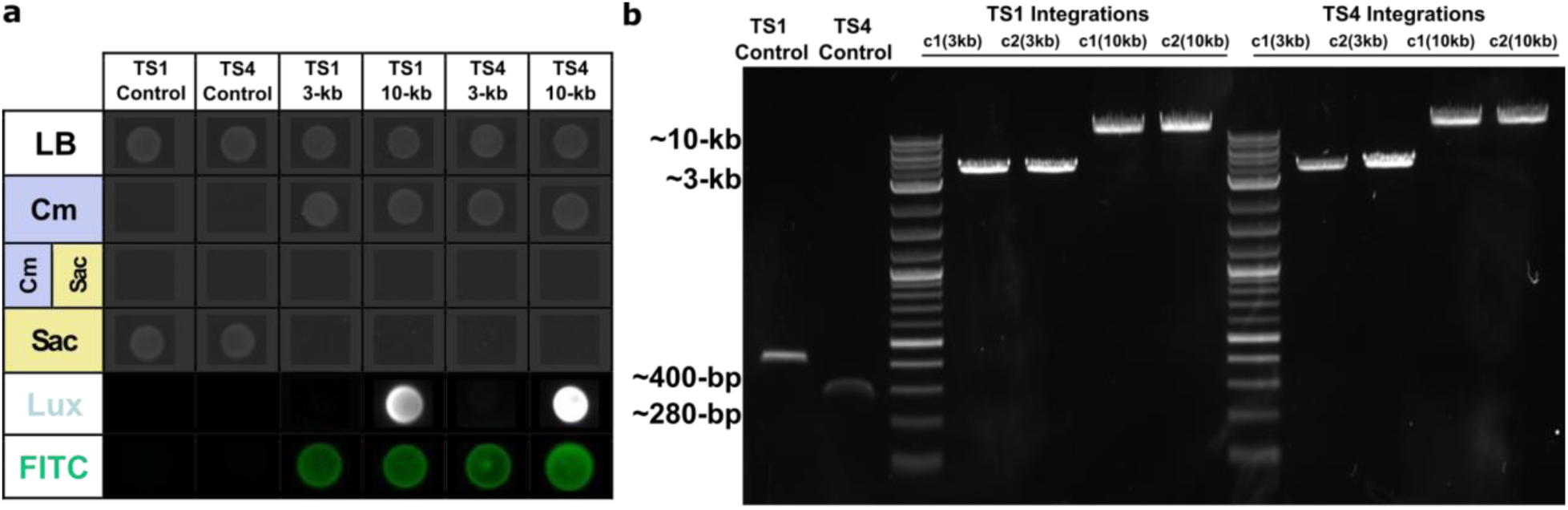
Phenotypic and genotypic screening comparing colonies from the 3-kb and 10-kb EM7-driven CAST integrations at TS1 or TS4. (A) Representative samples from phenotype spotting assays on selective plates; Cm, chloramphenicol; Sac, sucrose; Lux, luminescence; FITC, GFP. (C) Representative samples of the direct colony genotypes are shown after 3-kb or 10-kb integration.

All clones screened in this assay for both the 3-kb and 10-kb EM7-driven insert (8/8 each per target locus in Supporting Information, Figure S4) exhibited the expected phenotypes (Figure 5a). For all samples tested per insert, downstream genotyping with primers flanking each insertion locus yielded the expected lengths across both target loci (Figure 5b). Altogether, these results demonstrate that our PCR-based system for CAST-driven integrations does not suffer from a significant drop-off for large-sized integration products, as typically observed for other PCR-based integration systems.

### Decoupling the crRNA from Ara-pEffector

Although CAST systems have been shown to harbor the potential for iteration, no steps have been taken toward the simple and quick exchange of CAST crRNAs in cells. To accomplish this feat, the crRNA was decoupled from the Ara-pEffector and included on a separate vector containing a positive/negative selection cassette conferring resistance to Hygromycin B (+3, *HygroR*) and susceptibility to 4-Chloro-L-phenylalanine (−3, *pheS*^*T251A/A294G*^), a gene encoding for the mScarlet (*mScar*) red fluorescent protein, and a low-copy temperature-sensitive pSC101 backbone (Figure 6a). This modular crRNA construct, dubbed pTarget, expresses the crRNA with the constitutive J23119 promoter and can be easily removed/lost in cells by growing at high temperatures (≥37ºC) and/or counter-selecting using the negative marker drug in the backbone (4-Chloro-L-phenylalanine; herein Cl-phe).

**Figure 6.**
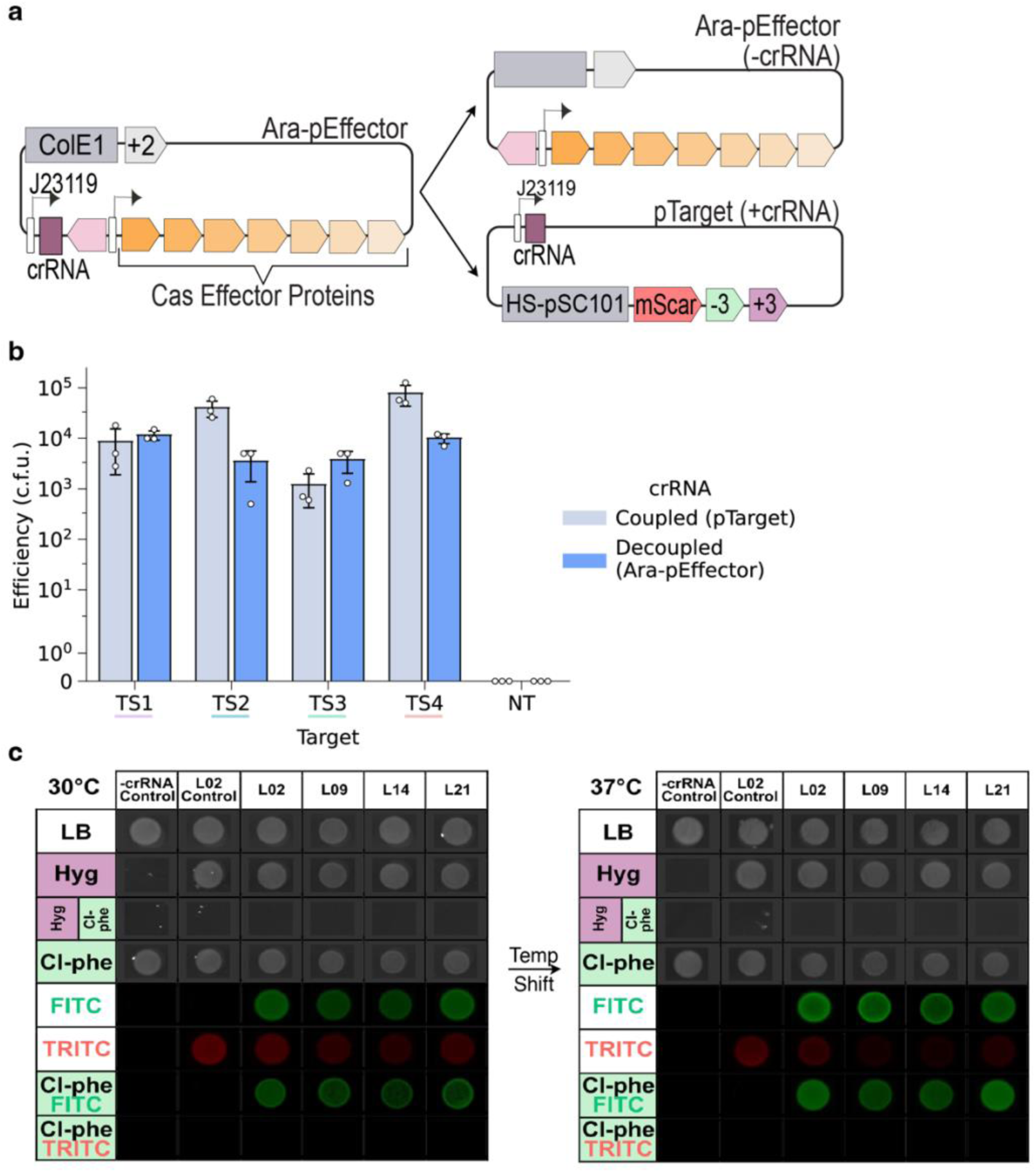
Schematic of pTarget construction and iterative insertion steps. (A) Creation of a two-plasmid Ara-pEffector and pTarget system that modularizes the targeting crRNA. (B) Integration efficiencies of 3-kb payload based on GFP+ CFU containing pTarget carrying crRNAs (biological replicates, n=3). (C) Representative colonies from the phenotype spotting assay were carried out at 30°C and 37°C for characterizing the removal of pTarget. Bars indicate the mean of biological triplicates with the error bars representing the standard error.

To test the functionality of the modular pTarget system, we transformed a 3-kb linear payload previously tested in this study into induced *E. coli* DH10b cells containing Ara-pEffector^-crRNA^ (without a crRNA) and a pTarget programmed to integrate at either TS1, TS2, TS3, or TS4. CFUs were characterized by the number of GFP+ colonies obtained following growth on selective agar plates. Based on this assay, the decoupled pTarget-pEffector system yielded similar integration efficiencies—on the order of 10^3^-10^4^ CFUs (Figure 6b)—as compared to results observed for the original, coupled system explored in Figure 3a. Our results, therefore, suggest that decreasing the copy number of the plasmid backbone carrying the crRNA does not affect our PCR-based CAST system’s ability to achieve high-efficiency genome integrations.

In order for the pTarget system to be viable for iterative genomic insertions, the pTarget plasmid must be rapidly lost on restrictive conditions. To test for the effectiveness of temperature and negative counter-selection on strains harboring pTargets, phenotyping assays were performed as described in previous experiments for 3-kb integrants. As noted, the positive/negative selection cassette included on the pTarget confers resistance to Hygromycin B and susceptibility to Cl-phe. Cells containing the pTarget plasmid will express mScarlet and therefore fluoresce under the corresponding excitation/emission wavelengths (Ex:569/Em:594). Phenotyping plates, all supplemented with Spectinomycin to maintain the Ara-pEffector^-crRNA^, were grown in permissive (30°C) and non-permissive (37°C) temperatures on varying selection compounds (Figure 6c). Red fluorescence intensity observed during microscopy of Cl-phe treated spots indicates that the higher, non-permissive temperatures only decrease the amount of pTarget plasmid present at the population level within a given spot. The absence of mScarlet fluorescence signal was only documented during fluorescence microscopy for cells spotted on Cl-phe, indicating that negative selection pressure is necessary and sufficient to fully select for plasmid loss. By coupling both heat sensitivity and negative selection mechanisms, we expected that our pTarget:pEffector CAST system would be capable of allowing us to quickly switch crRNA targets by iterative loss and transformation of pTargets.

### pTarget Plasmid System Enables Streamlined Iterative Insertions

To evaluate our design’s ability to modularly achieve and facilitate iterative insertions, we performed consecutive integrations in TS1 and TS4 (Figure 7a). A colony that had successfully integrated the 3-kb cargo into the TS1 site (verified through PCR based genotyping, as seen in Figure 7b) was re-streaked on agar plates containing spectinomycin on non-permissive temperatures (37 ºC) and Ch-phe to achieve the loss of the pTarget plasmid while maintaining the Ara-pEffector^-crRNA^. Loss of the pTarget plasmid was confirmed via red fluorescence microscopy and phenotyping assaying for resistance to Hygromycin B (not shown). Cells were then made electrocompetent prior to transforming a new pTarget harboring a crRNA targeting the TS4 site. Decreased efficiencies of transposition have been observed for target sites ≤1-Mb in distance to prior *V. cholerae* Type I-F CAST transposition events—a phenomenon generally referred to as transposon immunity^*50*^. Given this fact, the second target site (TS4) was chosen considering that it is ≥1-Mb away from the initial integration site. Following the acquisition of the new TS4-pTarget, cells were made electrocompetent under Ara-pEffector^-crRNA^ induction conditions. A new, ∼2-kb positive/negative selection cassette conferring kanamycin resistance (*KanR*) and streptomycin susceptibility (*rpsL*^*WT*^; nullifies the streptomycin resistance conferred by the DH10b *rpsL*^*K43R*^ genomic background) was transformed into the electrocompetent cells. Cells containing both the new 2-kb and previous 3-kb inserts were selected on agar plates containing both kanamycin (selecting for second insert) and chloramphenicol (selecting for first insert).

**Figure 7.**
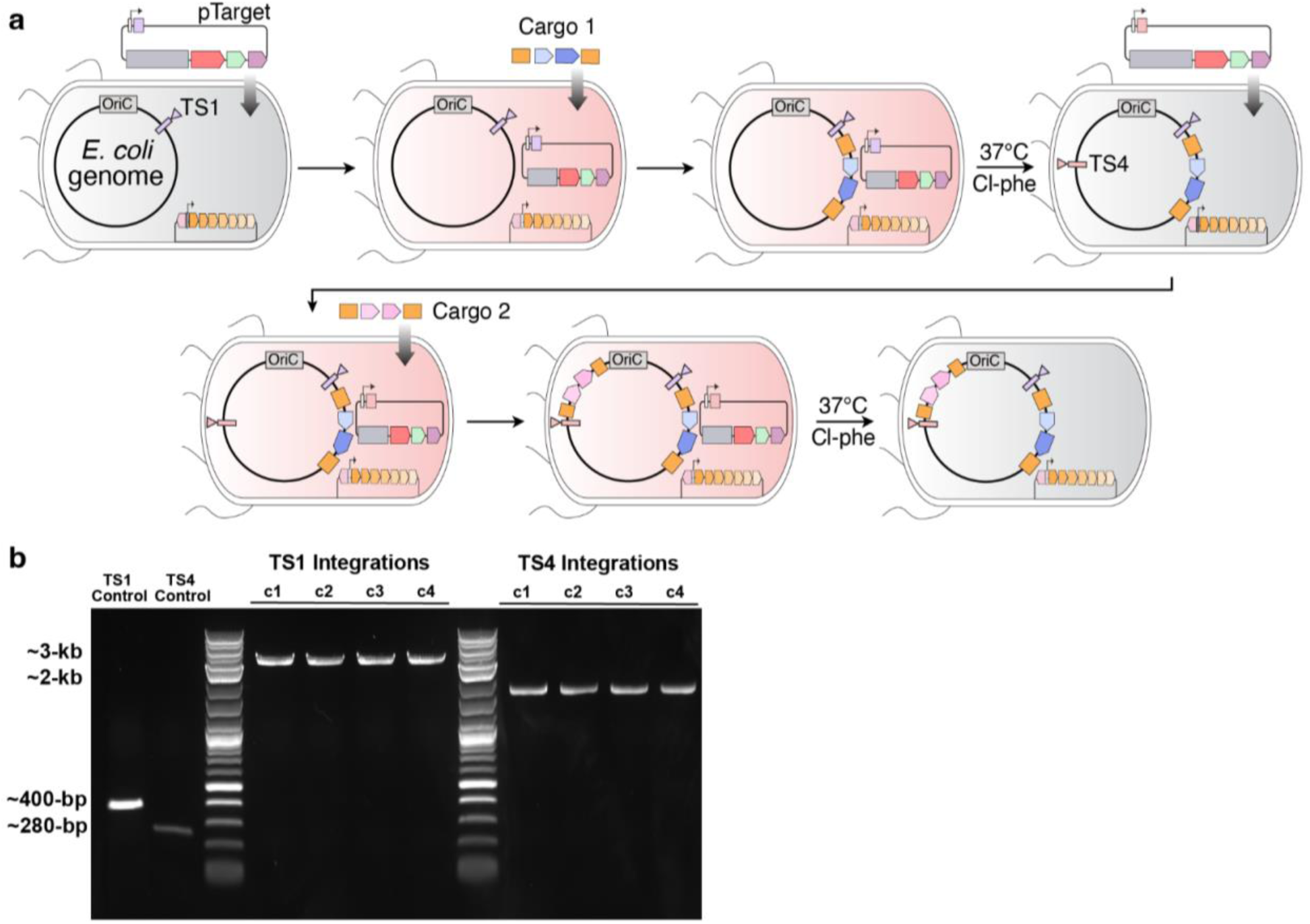
Iterative integrations with the removable pTarget plasmid. (A) Steps for iterative insertions in TS1 and TS4. Following transformation of the final transposable amplicon, the crRNA encoded within the pTarget guides transposition into the target site (here, TS1). The modular pTarget plasmid is then removed through growing at 37°C and on Cl-phe after the first insertion in TS1. The next pTarget containing a crRNA targeting TS4 is transformed before insertion of the second cargo into TS4. Finally, this pTarget is removed through the same growth conditions (37 ºC and Cl-Phe treatment). (B) Representative samples of the direct colony genotypes at both the TS1 and TS4 loci were conducted after iterative integration.

Following transformation of the first transposable amplicon, the crRNA encoded within the pTarget guides transposition into the first target site (TS4). The modular pTarget plasmid is then removed through growing at 37°C and on Cl-phe after the first insertion in TS1. The next pTarget containing a crRNA targeting TS4 is transformed before insertion of the second cargo into TS4. Finally, this pTarget is removed through the same growth conditions (37 ºC and Cl-Phe treatment). Genotyping of the two junctions (TS1 and TS4) revealed that both the 3-kb and 2-kb payloads were respectively present on the correct genomic loci (Figure 7b). These results confirm that pTarget exchange allows for iterative insertions of different payloads in a seamless manner, expanding on the programmability of CAST-based genome integration systems in bacteria.

## Conclusion

Recently discovered CRISPR-associated transposons (CASTs) have been used to genomically integrate DNA payloads in bacteria. Although these systems show promise in the realm of genome engineering, the current implementation of CAST-based integration systems lacks modularity and streamlining capabilities. In order to expedite and streamline genomic integration using CASTs, we bypassed the need to construct intermediate plasmids harboring the desired cargo insert by reconstituting the L and R transposon arms through a two-step, nested PCR. In our experiments, site-specific transposition of linear cargo products produced through this approach was shown to be agnostic to sequence length, identity, and genomic target. Direct comparison with the Lambda-Red recombination system demonstrated that CAST systems can integrate tested PCR amplicons with orders of magnitude higher efficiency than the Lambda-Red system, enabling the integration of amplicons with comparable efficiencies across distinct genomic loci. To increase the versatility of the system, we decoupled the targeting crRNA from the CAST-Effector plasmid by placing the crRNA into a removable targeting plasmid, which we called pTarget. The use of the pTarget plasmid alongside Ara-pEffector^-crRNA^ plasmid yielded high amplicon integration efficiencies (∼10^4^ CFU), and allowed for iterative CAST genomic insertions after exchange of pTarget-crRNA constructs targeting distinct genomic loci. These results may be translated to other CAST systems, expediting the process in which future CAST systems are experimentally screened for and tested. Altogether, our CAST-directed nested PCR method enables efficient and precise genome integrations without the need for preparatory cloning, and implementation of this system could further benefit the streamlined engineering of more industrially-relevant microorganisms, such as *Pseudomonas putida*.

## Materials and Methods

### Bacterial Strains and Growth Conditions

Experiments and plasmid construction were conducted in *E. coli* DH10b (Invitrogen), which carries an K43R mutation in *rpsL* that confers resistance to streptomycin. Cells were cultured at 30°C or 37°C (170 rpm) in lysogeny broth (LB medium) (1% (w/v) tryptone, 0.5% (w/v) yeast extract, 1% (w/v) NaCl). As needed, selective LB cultures were prepared with chloramphenicol (25 μg/mL^−1^), spectinomycin (50 μg/mL^−1^), hygromycin (200 μg/mL^−1^), kanamycin (50 μg/mL^−1^), sucrose (7.5% w/v), and/or 4-chloro-phenylalanine (2.5 mM). LB agar plates were prepared by supplementing LB medium with 1.5% (w/v) agar.

### Plasmid and Nested PCR Construction

Names and descriptions of plasmids and their sources are listed in Table S1. Genbank annotation files for all plasmids used in this study are deposited here: https://github.com/meganwang08/A-universal-system-for-streamlined-genome-integrations-with-CRISPR-associated-transposases. Ara-pEffectors were constructed from pSL1777 (Addgene #160731) in three separate pieces through Gibson assembly: the pCDF backbone containing SmR and CAST effector proteins, the J23119 promoter with the targeting crRNA, and araC with the pBad promoter. Subsequent targeting Ara-pEffectors (TS1, TS2, TS3 and TS4) were subcloned from this construct by digesting with PmlI and XhoI restriction enzymes (NEB) and performing a two-piece Gibson assembly with the digested fragment and the crRNA fragment. crRNA fragments were generated through PCR amplification of two primers encoding the crRNA and Gibson homology sequences without a template. -crRNA Ara-pEffector was built by PCR amplifying the CAST proteins from the original Ara-pEffector and the pCDF backbone containing SmR and assembling in a two-piece Gibson. pTarget constructs were built via two-piece Gibson assembly. The first piece contained the heat-sensitive backbone, positive/negative selection cassette conferring hygromycin resistance and Cl-phe susceptibility, and mScarlet. The second piece contained the respective crRNA driven by the J23119 promoter. The Lambda-Red recombinase plasmid, pKW20, was taken from previous work^17^. Primers used to clone all plasmids are listed in Table S2. Gibson assemblies were performed as described in the HiFi DNA Assembly protocol by New England Biolabs. Nested PCR primers were ordered from Sigma-Aldrich’s custom oligo service. Oligo design and the sequences used for the reconstitution of the R and L transposon arms are provided in Figure 1b. As illustrated in Figure 1b, the primer pair for the first round of the nested PCR (Primer 1: 5’-NNNNN(cargo)-3’, Primer 2: 5’-CCACAAAACAACCATATATTGATATCTCACAAAACAACCATAAGTTGATATTTTTGTGAATCGAGTATTTCAGCAAAACTACTGCAGTAAG-NNNNN(cargo)-3’) cargo binding only and partially reconstituted L arm. The primer pairs for the second round of the nested PCR (Primer 1: 5’-TGTTGATACAACCATAAAATGATAATTACACCCATAAATTGATAATTATCACACCCA-NNNNN(cargo)-3’, Primer 2: 5’-TGTTGATGCAACCATAAAGTGATATTTAATAATTATTTATAATCAGCAACTTAACCACAAAACAACCATATATTGATATCTCACAAAACAAC-3’) fully reconstituted both R and L transposon arms. PCRs used the PrimeSTAR GXL DNA Polymerase (Takara Bio) under conditions following the protocol listed by the supplier. In summary, all PCRs were set up for 36 cycles using the following settings: 1 min at 98°C for the initial denaturing step, 10 seconds at 98°C for cycle denaturing step, 15 seconds at 60°C for cycle annealing step, 1 minute per kilobase at 68°C for cycle elongation step, 5 minutes at 68°C for a hold step. The amplicon product from the first PCR round was diluted to approximately 5 ng/μL to use as template for the second nested PCR round. The exact primer sequences used in the nested PCRs performed in this work are listed in Table S2.

### Electrocompetency, Chemicompetency, and Transformation Procedures

Electrocompetent cells were prepared for experiments performed in this work. 0.5 L of cells were made electrocompetent and induced at OD600 0.2 with 0.5% L-arabinose for 1 hour when necessary. Upon reaching OD600 0.5, cultures were chilled on ice for 30 minutes. Three subsequent washes at 0°C 4000 rpm were performed: two rounds resuspending the pellet with MilliQ (Sigma-Millipore) ddH2O and one round resuspending the pellet with pre-autoclaved 15% glycerol. Cells were resuspended in 500 μL of pre-autoclaved 15% glycerol and 110 μL was aliquoted into 1.5mL tubes and stocked by flash freezing in liquid nitrogen. An Eppendorf Eporator was used to transform electrocompetent cells, using a 2500V setting and 2-mm gap electroporation cuvettes. Electroporated cells were quickly recovered in 1 mL of LB for 1 hour (900 orbital rpm) at 30°C or 37°C, depending on the temperature tolerance of the transformed vector backbone(s).

Chemicompetent cells were prepared to transform sequence-verified pTarget plasmids into Ara-pEffector(−crRNA) cells in the process of strain building. Chemicompetent cells were prepared in the same growth conditions at 10mL but resuspended with 250 μL TSS (10% w/v PEG-3350, 5% w/v DMSO, 20mM MgCl_2_) and 50 μL KCM(0.5M KCl, 0.15M CaC_l2_, 20mM MgCl_2_) after centrifugation at 0°C 4000 rpm. Heat shock (4°C for 10 minutes, 42°C for 90 seconds, and 4°C for 3 minutes) was used to transform sequence-verified pTarget plasmids into 50 μL of chemicompetent cells and these cells were recovered in 150 μL of LB at 30°C for 1 hour.

### CAST-directed Integrations and Lambda-Red Recombinations

For CAST-directed integrations, 200 μL of nested PCRs were purified as specified by the Qiagen QIAquick PCR Purification Kit. Importantly, the purified nested PCR product was immediately applied to downstream CAST-directed integrations as freeze-thaw of the linear DNA led to inconsistencies in reported integration efficiencies. 1.2 μg of DNA product was transformed into induced electrocompetent cells at 2500V containing the respective plasmid(s) (either Ara-pEffector or Ara-pEffector(−crRNA) with pTarget). Transformants were recovered in 4 mL LB at 30°C for four hours with further induction with 80 μL of L-arabinose (0.5%) at the one-hour mark. The 4 mL culture was further diluted with 10 mL LB containing antibiotics selecting for both the plasmids present and the transformed cargo. The selective LB culture was recovered overnight for 18 hours at 30°C. For spreading cells on selective LB agar, four dilutions were used for plating: 180 μL of the culture was taken as the neat condition while 180 μL of a 1:10, 1:100, or 1:1000 dilution in ddH_2_O was spread for the diluted conditions. Finally, the plates were incubated at 30°C for 22 hours and transferred to 37°C for four more hours before colonies were counted. GFP+ and bioluminescence were quantified with the ChemiDoc Imaging System from BioRad using the Blue Tray SYBR Safe (0.5 seconds exposure) and Chemi/UV/Stain-Free Tray chemiluminescence (30 seconds exposure) settings, respectively. CFUs were calculated from only the fraction of the culture that was spread on plates.

PCR products for Lambda-Red integrations were DpnI digested overnight to remove background template and purified as specified by the Qiagen QIAquick PCR Purification Kit. Following purification, 1.2 μg of DNA product containing 50-bp homology on both the 5’ and 3’ ends to the TS1 and TS4 integration sites were transformed into induced cells containing a plasmid with Lambda-Red (pKW20)^17^. The transformants were recovered in 4 mL of LB at 30°C for four hours. No additional induction step occurred at the 1 hour since excess Gamma protein has been shown to be toxic to cells. The 4 mL culture was further diluted with 10 mL LB containing antibiotics selecting the transformed cargo. The selective LB culture was recovered overnight for 18 hours at 30°C. The recovered cells were spread on selective plates as described for CAST-directed integrations. Plates were incubated at 37°C until colonies formed and CFUs were counted as described for CAST-directed integrations.

### Population Next-Generation Sequencing (NGS) Preparation and Data Analysis

Approximately 8000 colonies were washed from CAST-directed integration experiment plates and cultured in 4 mL selective LB. 1 mL of the culture was aliquoted for genomic DNA extraction using the Qiagen DNeasy Blood & Tissue Kit. 5 ng of the extraction for each genomic target site (TS1, TS2, TS3, TS4) was amplified with the PrimeSTAR GXL DNA Polymerase in 50 μL reactions each. Amplification primers were designed to bind ∼200-bp away from each integration site to capture the insertion junction. Reactions were purified with the Qiagen QIAquick PCR Purification Kit and sequenced through core facilities at the California Institute of Technology (Millard and Muriel Jacobs Genetics and Genomics Laboratory). Samples were sequenced on an Illumina HiSeq instrument using a 50-bp paired-end chemical reagent kit.

To distinguish, quantify, and analyze on-target insertions, alignment reference sequences were generated for genomic loci assessed in this work assuming an integration range between 40-60 bp away from the genomic target site in both RL and LR directions. Reference sequences were compiled in a multi-FASTA reference file, and the Burrows-Wheeler Alignment (BWA)^51^ tool’s BWA-MEM setting was used to align Illumina reads against the indexed multi-FASTA file. The alignment output (SAM) was converted to BAM format, sorted, and indexed using SAMtools^52^. Mapped reads were extracted into a raw FASTQ file. Aligned reads were then further analyzed for on-target integrations, directionality, and integration location. Read coverage was graphed by generating a BedGraph with bamCoverage from the deepTools package^53^.

On-target and off-target integration events were quantified by using a 20-bp pre-integration junction created from the 40-60-bp downstream of the target site as a sliding window to perform a substring search against aligned reads. The total number reads matching the pre-integration junction was divided by the total number of initial aligned reads to calculate an off-target percentage. On-target percentages were calculated by taking the complement of the off-target percentage. The directionality of the insert (RL or LR) and position of insertion was given by the total number of reads that matched each reference in the multi-FASTA file mentioned above. For directionality, percentages were calculated by counting reads that aligned to either the RL or LR orientation reference files and dividing by the total number of aligned reads. For the position of insertion, percentages were calculated by dividing the number of reads aligned to each multi-FASTA file by the total number of reads aligned per direction (RL or LR). Pipelines generated to analyze sequencing data were written in Python and are publicly available in a GitHub repository. Raw sequencing data used for analysis in this work can be found under NCBI’s Sequence Read Archive (PRJNA835982).

### Phenotyping Spotting Assays and Genotyping PCR Analyses of Transposition

Single colonies from transformation plates were picked and resuspended in 60 μL ddH_2_O and 4 μL of the resuspension were taken to spot on selective plates for phenotyping. Plates were incubated at 30°C or 37°C depending on the strain and conditions for 18 hours. The resuspensions were then used for PCR amplification across the genomic region of the expected insertion junction. Primers were designed to anneal to the 200-bp flanking the insertion junction. PCRs were performed using PrimeSTAR GXL DNA Polymerase according to the supplier protocol. 1 uL of resuspended cells were directly added to 19 uL PCR reaction and lysed at 98°C for two minutes prior to the PCR cycles as described above. PCR amplification was run for 28 cycles rather than the usual 36 cycles to avoid non-specific amplification. 2 uL of PCR products were mixed with 5.5 uL ddH_2_O and 1.9 uL 6X gel loading dye (NEB) and loaded onto 1% w/v agarose gels cast with SYBR Safe DNA gel stain. Gels ran at 100V for 30 minutes before imaging with the ChemiDoc Imaging System using the Black Tray SYBR Safe option (optimal auto-exposure setting).

### Fluorescence Microscopy and Bioluminescence Imaging

Fluorescence from GFP and mScarlet reporters on the phenotyping plates was assessed with an Olympus Apochromat N 1.25X/0.04 na objective on an Echo Revolve microscope in the upright position setting. Exposure for the GFP and mScarlet channels was set to 70 ms and 50 ms, respectively. Light intensity was set to 45% for GFP and 71% for mScarlet channels, and images were taken with a low gain setting. Images were pseudocolored through built-in Echo Revolve software based on signal intensity and the expected emission wavelength for the measured fluorophore signal. Bioluminescence from phenotyping spots was assessed on a ChemiDoc Imaging System (BioRad) using the Chemi/UV/Stain-Free Tray chemiluminescence (30 seconds exposure) settings. To eliminate bioluminescent signals from other integration location queries during the imaging process, colonies from other transformations were covered to provide the cropped phenotype images.

## Supporting Information

- Phenotype spotting assays and genotype gels of Ara-pEffector TS1, TS2, TS3, TS4 integrations (Figure S1).
- Sanger sequencing chromatagrams for 3-kb TS1, TS2, TS3, and TS4 (Figure S2).
- Select maps of read coverage from Illumina amplicon sequencing across genomic inserts (Figure S3).
- Genotype of the junctions containing 3-kb or 10-kb Em7-driven insertions (Figure S4).
- Ara-pEffector efficiencies (Figure S5) and respective phenotype spotting (Figure S6) for experiments in Figure 3.
- Gel of amplicons used in NGS sequencing (Figure S7).
- Ara-pEffector efficiencies (Figure S8) and Lambda-Red efficiencies (Figure S9) for J23119-driven cargos in Figure 4c.
- Genotypes of Lambda-Red integrations (Figure S10).
- Phenotype spotting for 3-kb and 10-kb EM7-driven integrations (Figure S11) for Figure 5a.
- pTarget efficiencies (Figure S12), respective phenotype spotting (Figure S13) for experiments in Figure 6.
- Genotype of both junctions (Figure S14) for iterative insertion experiments in Figure 7.
- Plasmid map of Ara-pEffector-TS1 (Figure S15) and pTarget-TS1 (Figure S16).
- Annotated plasmid sequence of Ara-pEffector-TS1 (Figure S17) and pTarget-TS1 (Figure S18).
- Table of plasmids used in this study (Table S1)
- Table of primers used in this study (Table S2)
- Table of genomic target site equivalencies (Table S3)

## Author Contributions

M.W. and C.S. conceived and designed this research. R.J.Z and M.W. cloned the plasmids used in this study. M.W. performed laboratory experiments and analyzed data. M.W. performed phenotyping analyses and R.J.Z. performed PCR-based genotype analyses. M.W. analyzed and graphed NGS data. M.W., C.S., and R.J.Z. wrote the paper with input from all authors. K.W. provided insightful guidance for the work.

### Funding

This work was supported by the Shurl and Kay Curci Foundation (grant no. 108625) and the NIH (DP2-GM140937).

### Notes

The authors declare no competing financial interest.

Data availability: we declare that the data supporting the findings of this study are available within the paper and its Supporting Information. NGS analysis pipelines can be found at https://github.com/meganwang08/A-universal-system-for-streamlined-genome-integrations-with-CRISPR-associated-transposases. Raw sequencing data produced and used in this study are available from the National Center for Biotechnology Information (NCBI) under SRA accession no. PRJNA835982.

## Acknowledgments

Work was supported by the Shurl and Kay Curci Foundation (grant no. 108625, K.W.) and the NIH (DP2-GM140937, K.W.). R.J.Z is supported by the National Science Foundation Graduate Research Fellowship (grant no. 1745301). We thank Igor Antoshechkin from the Millard and Muriel Jacobs Genetics and Genomics Laboratory for sequencing library preparations and Illumina amplicon NGS.

## Notes

https://github.com/meganwang08/A-universal-system-for-streamlined-genome-integrations-with-CRISPR-associated-transposases

https://www.ncbi.nlm.nih.gov/sra/PRJNA835982

### Competing Interest Statement

The authors have declared no competing interest.

