## Supplemental for "A universal system for streamlined genome integrations with CRISPR-associated transposases"

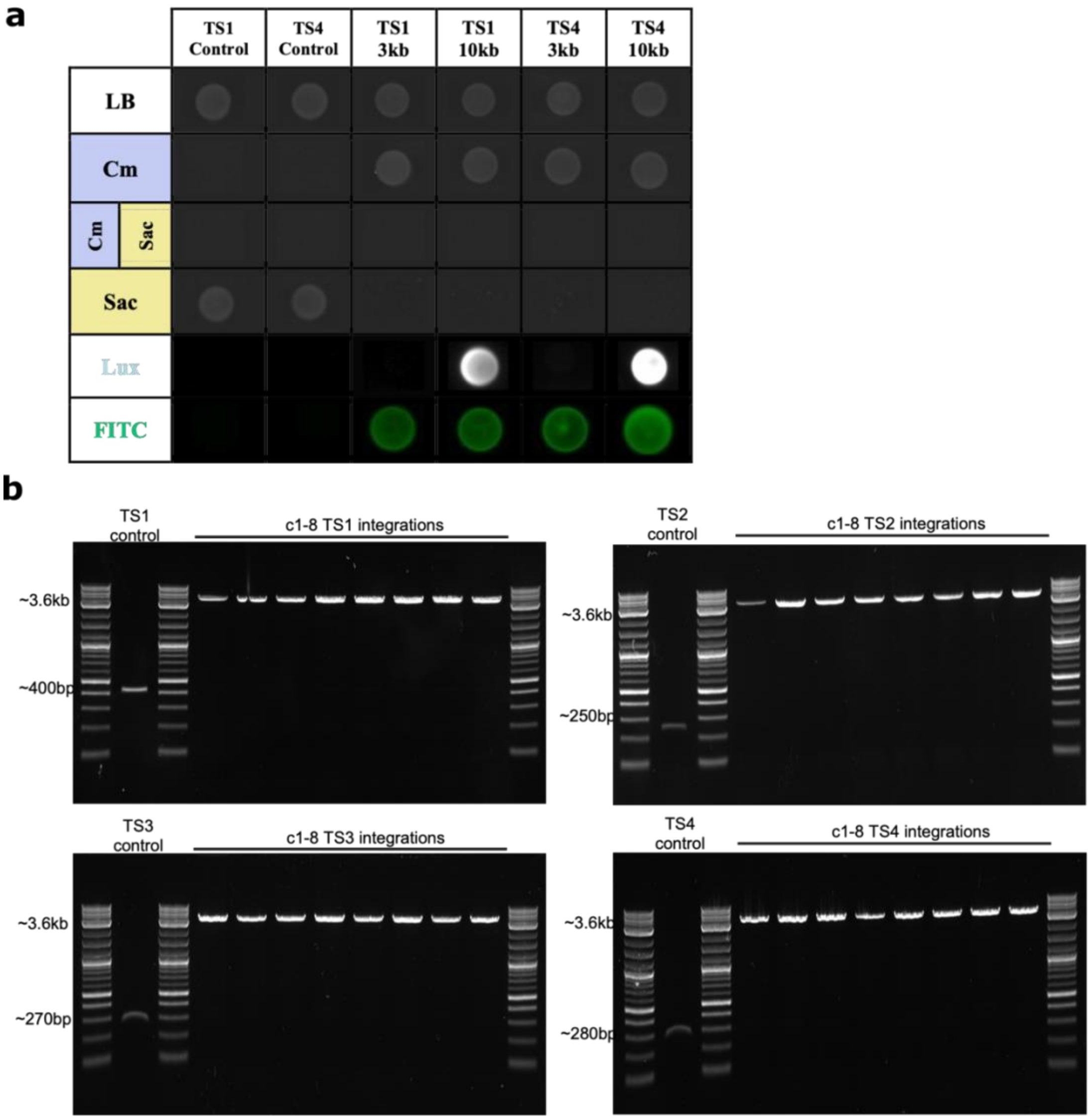


Figure S1. Phenotypic and genotypic assays of Ara-pEffector TS1, TS2, TS3, TS4 integrations. (A) Representative phenotype spotting assays on selective agar plates. (B) Genotype of the insertion location after Ara-pEffector mediated integration of a 3-kb payload.

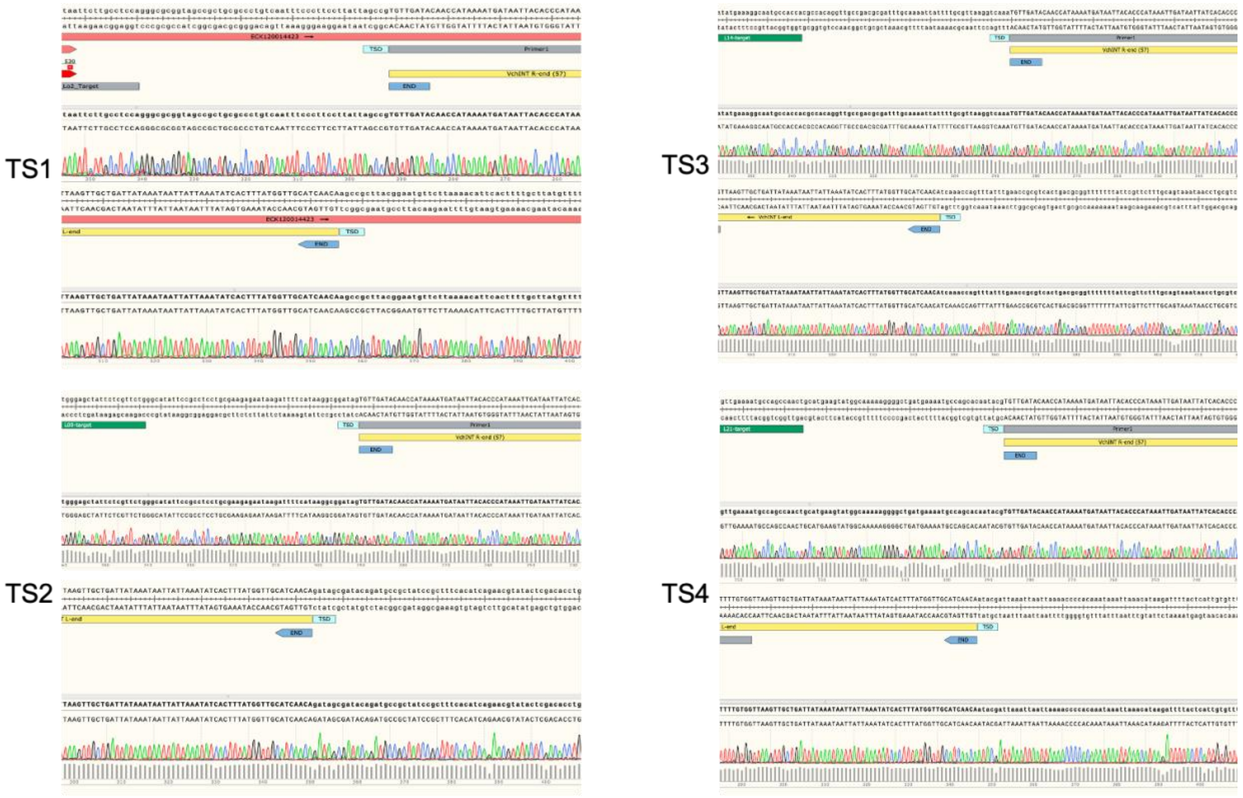


Figure S2. Sanger sequencing chromatograms of genome junction post-integration of 3-kb cargo at TS1, TS2, TS3, and TS4. The target site duplication (TSD) is annotated upstream and downstream of the integration sites.

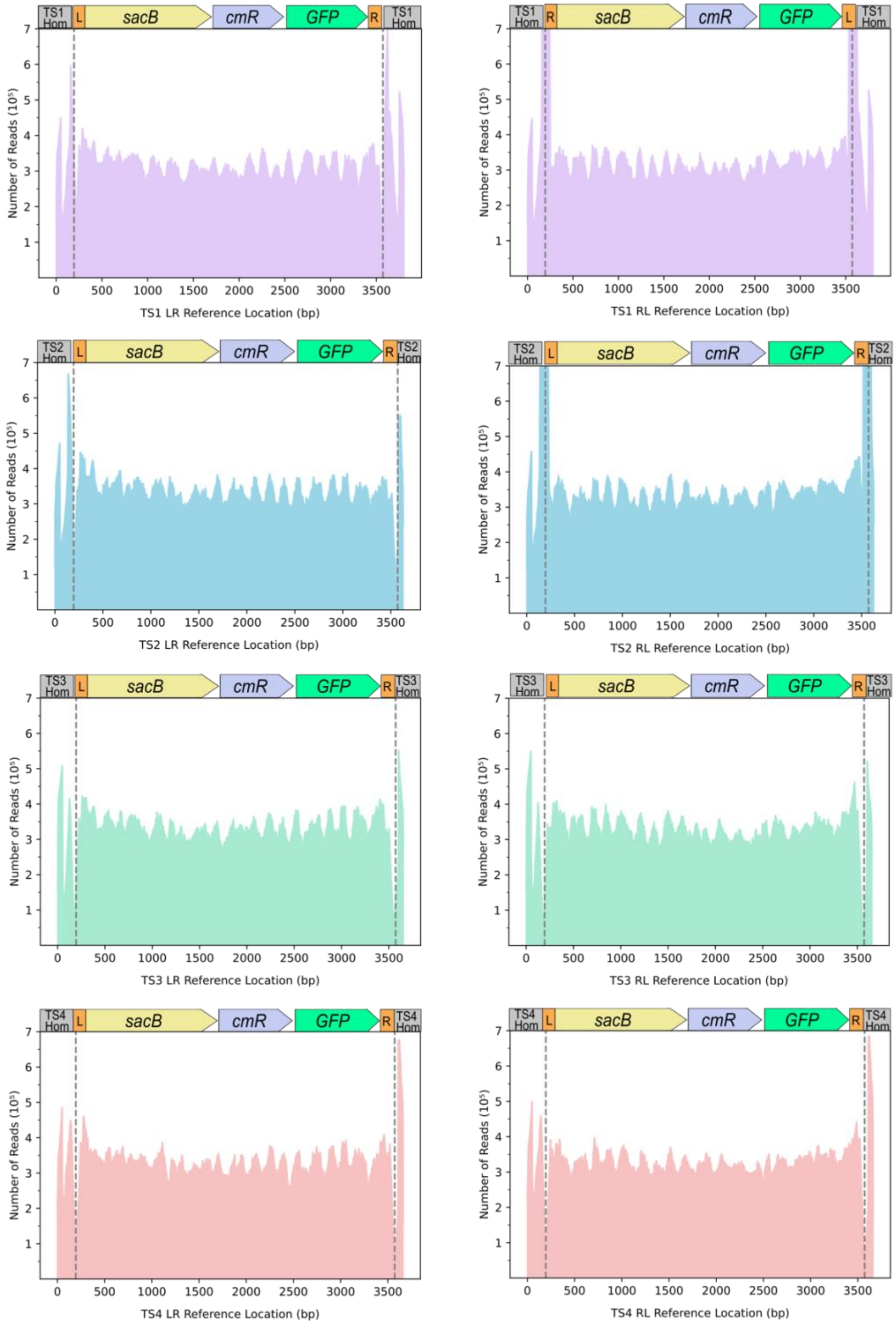


Figure S3. Select maps of read coverage from Illumina amplicon sequencing across genomic inserts. The alignment files of read coverage were chosen based on the most highly integrated position for each genomic location and direction given in figure 2d.

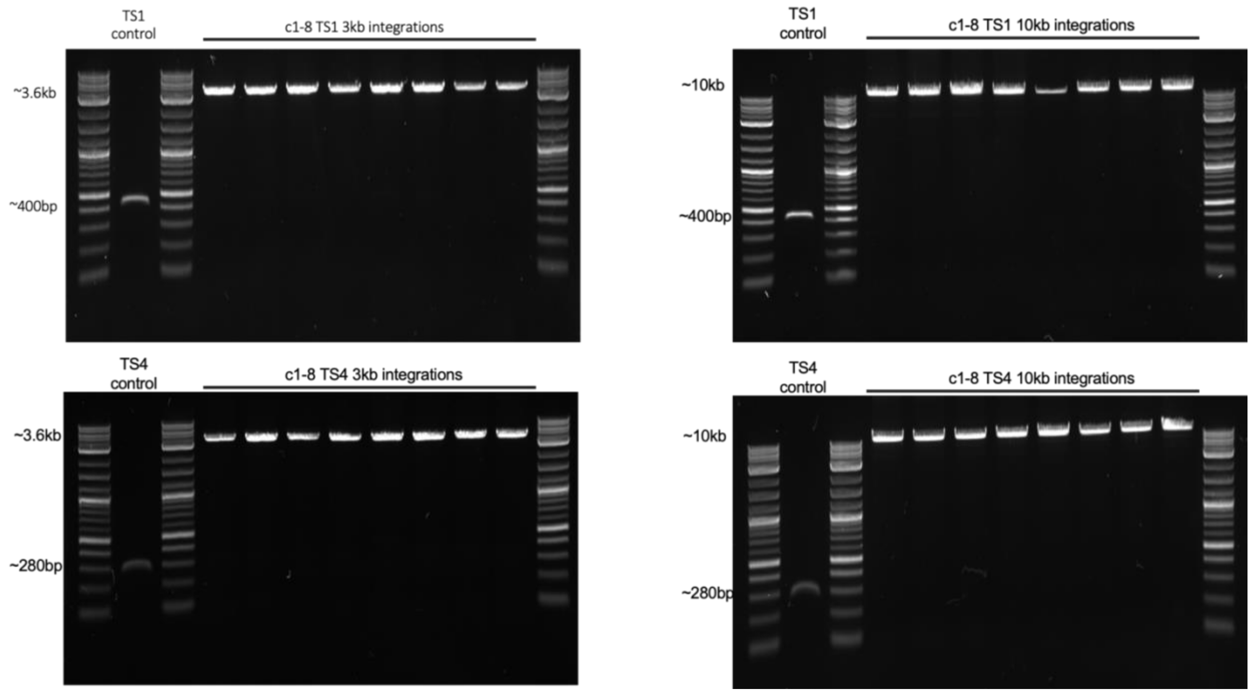


Figure S4. Genotype of the junctions containing 3-kb or 10-kb insertions in either the TS1 or TS4 position.

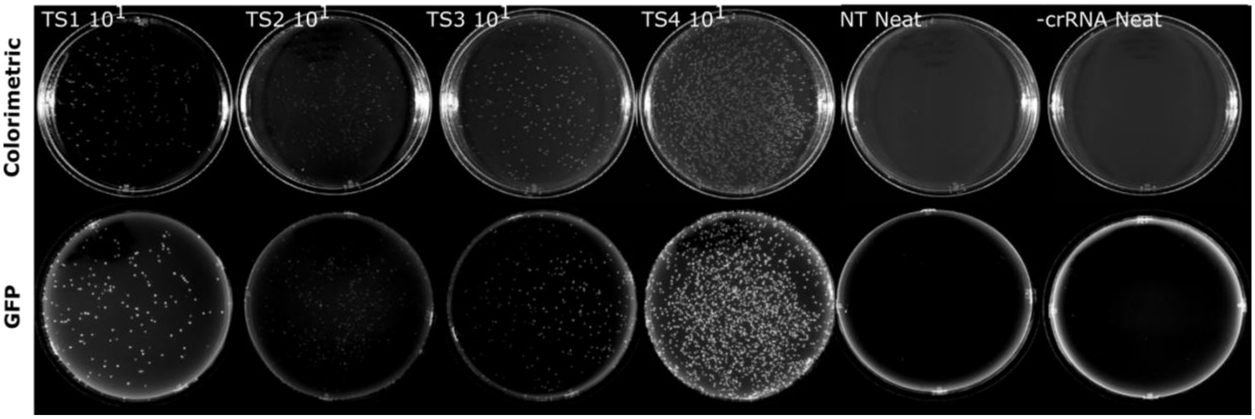


Figure S5. Representative colorimetric (top) and GFP images (bottom) of Ara-pEffector efficiency experiments (see Figure 3a in the main text) on $10^1$ or neat dilution plates for various crRNAs (TS1, TS2, TS3, TS4, NT, -crRNA).

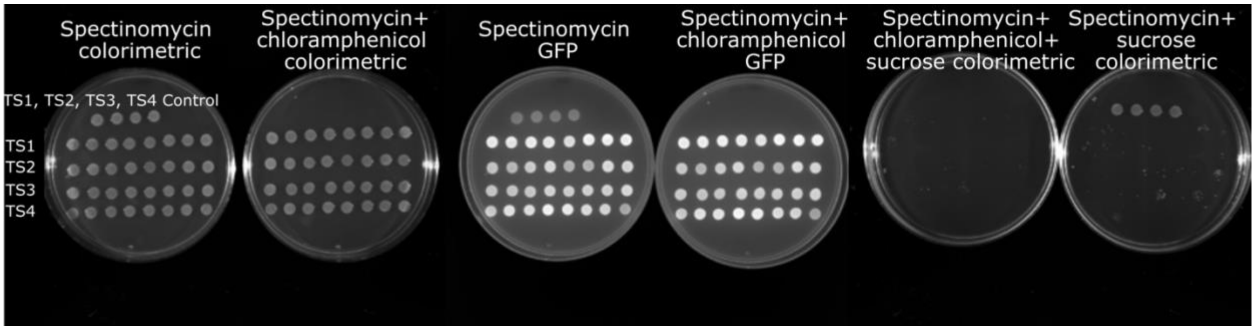


Figure S6. Phenotype spotting plates for TS1, TS2, TS3, TS4 (8 colonies each) with the four respective pre-integration Ara-pEffector controls at the top.

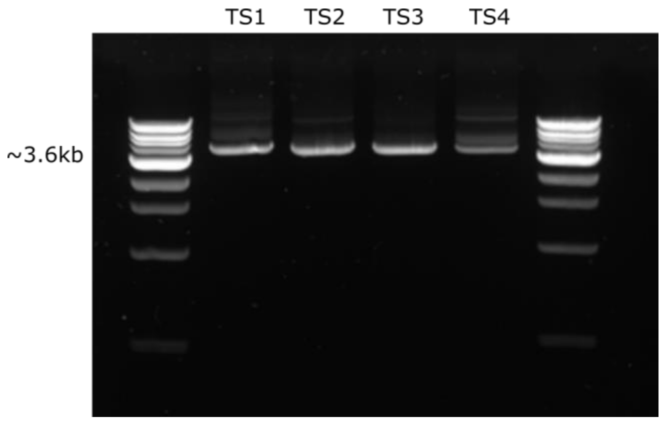


Figure S7. Amplicons used in NGS sequencing.

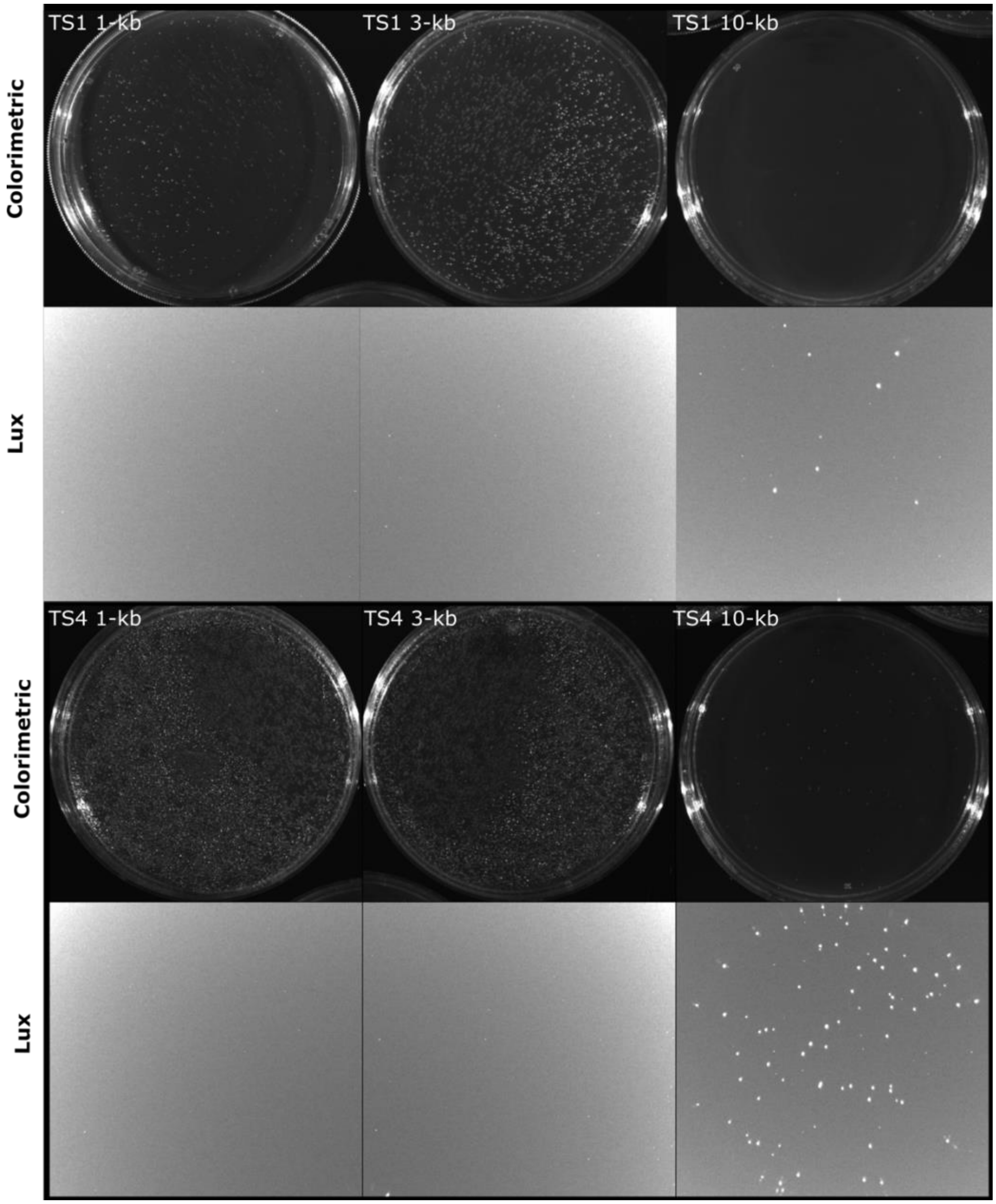


Figure S8. Representative colorimetric and bioluminescence images of Ara-pEffector 1-kb, 3-kb, and 10-kb CAST integration efficiency experiments (see Figure 4c in the main text) on the $10^1$ dilution plate at TS1 and TS4. These experiments were carried out in parallel with Lambda Red 1-kb, 3-kb, and 10-kb integrations at TS1 and TS4 (Figure S9).

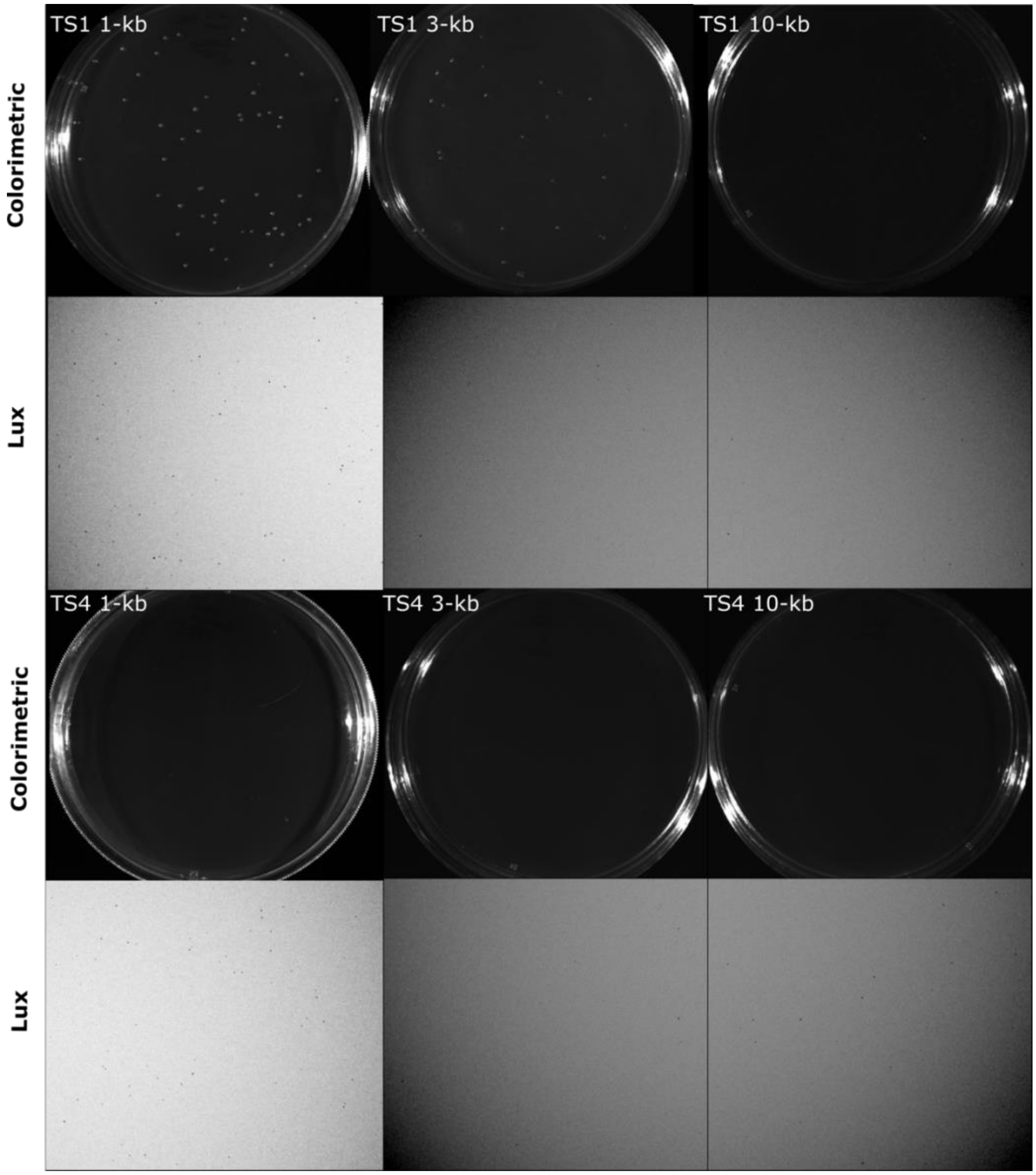


Figure S9. Representative colorimetric and bioluminescence images of 1-kb, 3-kb, and 10-kb Lambda-Red integration efficiency experiments (see Figure 4c in the main text) on the neat dilution plate at TS1 and TS4. These experiments were carried out in parallel with CAST 1-kb, 3-kb, and 10-kb integrations at TS1 and TS4 (Figure S8). Colonies were screened against background template through the absence of bioluminescence in the 1-kb and 3-kb integration colonies.

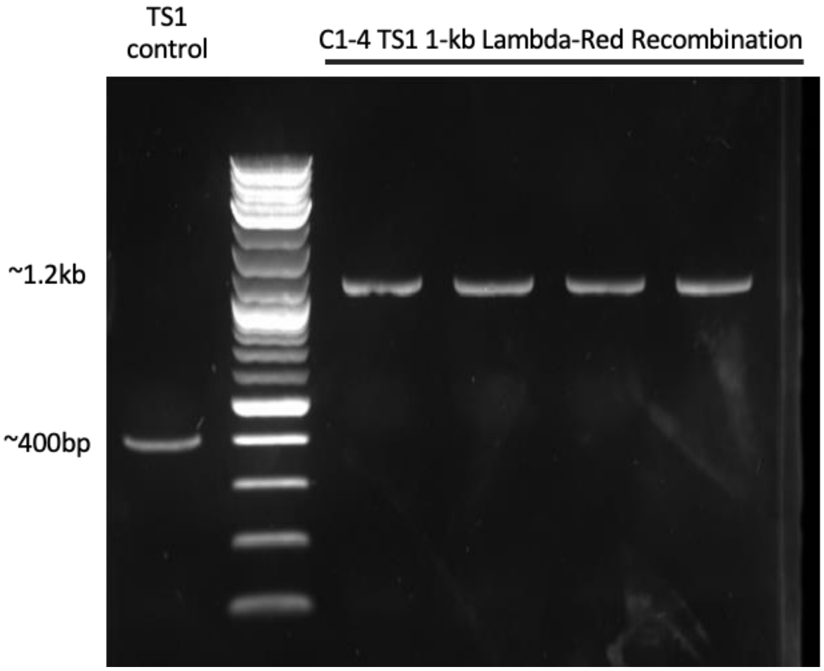


Figure S10. Genotypes of four colonies for Lambda Red mediated recombination of 1-kb cargo in the TS1 site (see Figure 4c in the main text).

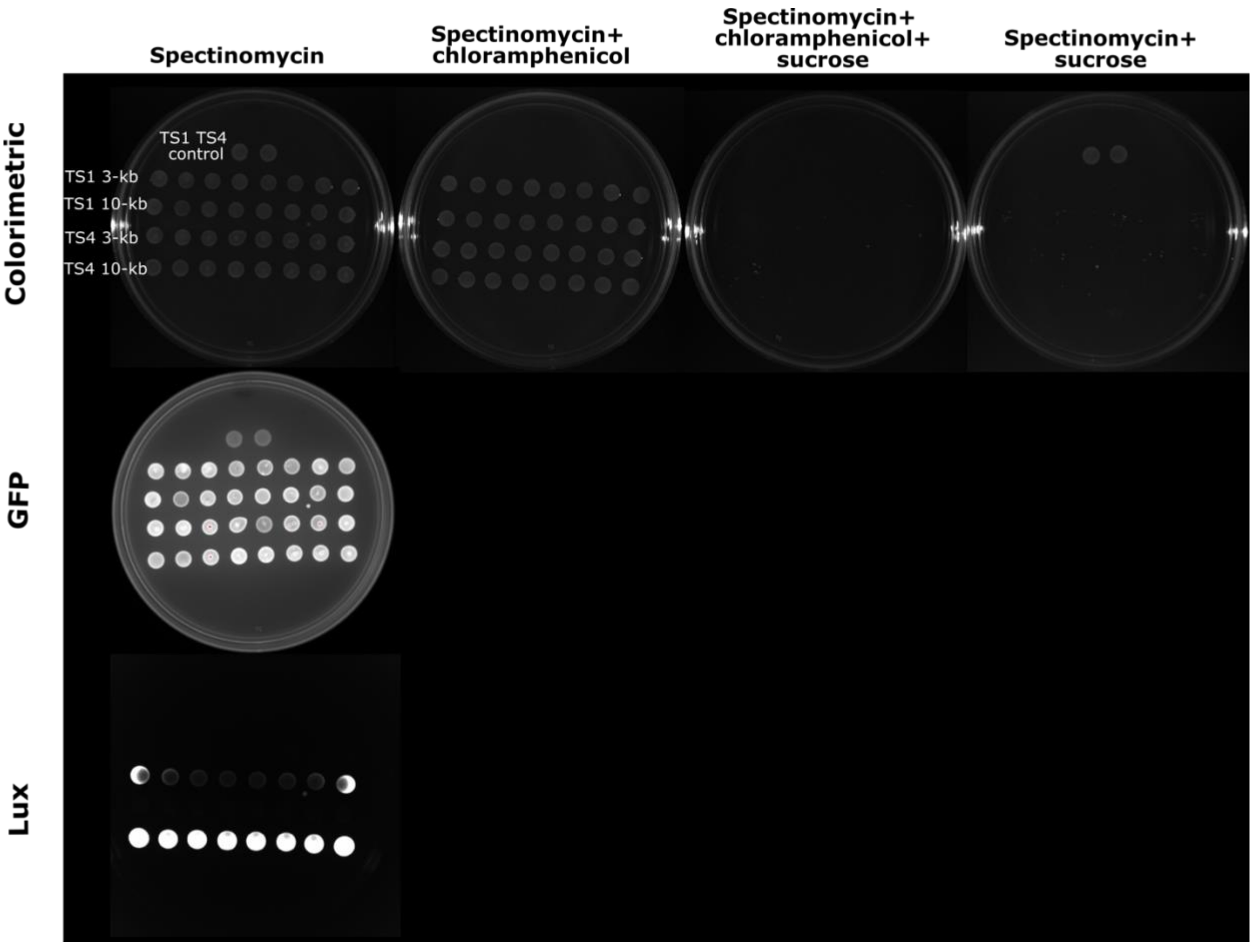


Figure S11. Phenotype spotting plates for Ara-pEffector with varying crRNAs (TS1 or TS4) and either the 3-kb or 10-kb insert.

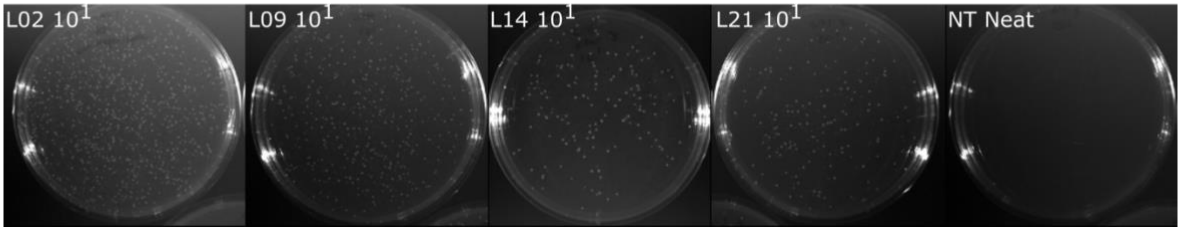


Figure S12. Representative colorimetric images of Ara-pEffector$^{-crRNA}$ with pTarget efficiency experiments (see Figure 7a in the main text) on $10^1$ or neat dilution plates for various crRNAs (TS1, TS2, TS3, TS4, NT).

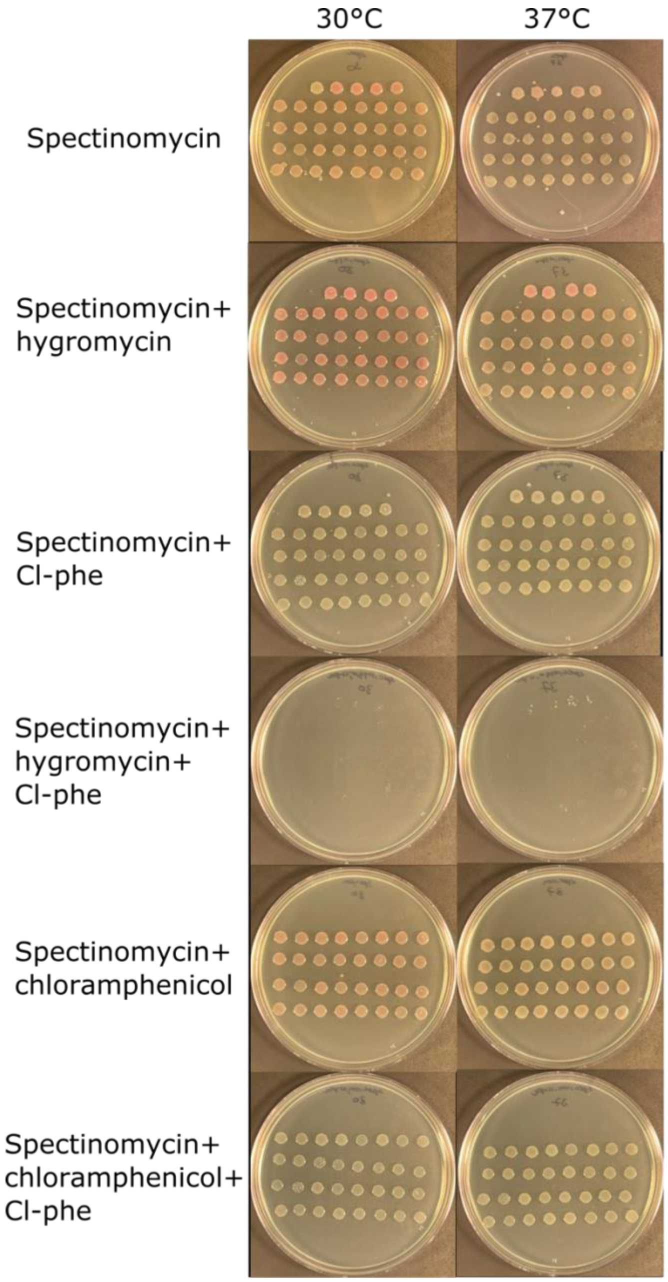


Figure S13. Phenotype spotting plates for Ara-pEffector$^{-crRNA}$ with pTarget TS1, TS2, TS3, or TS4 integrations of 3-kb insert. Top row per plate shows controls (from left to right): Ara-pEffector$^{-crRNA}$, pre-integration Ara-pEffector$^{-crRNA}$ with pTarget TS1, pre-integration Ara-pEffector$^{-crRNA}$ with pTarget TS2, pre-integration Ara-pEffector$^{-crRNA}$ with pTarget TS3, and pre-integration Ara-pEffector$^{-crRNA}$ with pTarget TS4. Eight post-integration colony replicates for TS1, TS2, TS3, and TS4 follow in the next four rows.

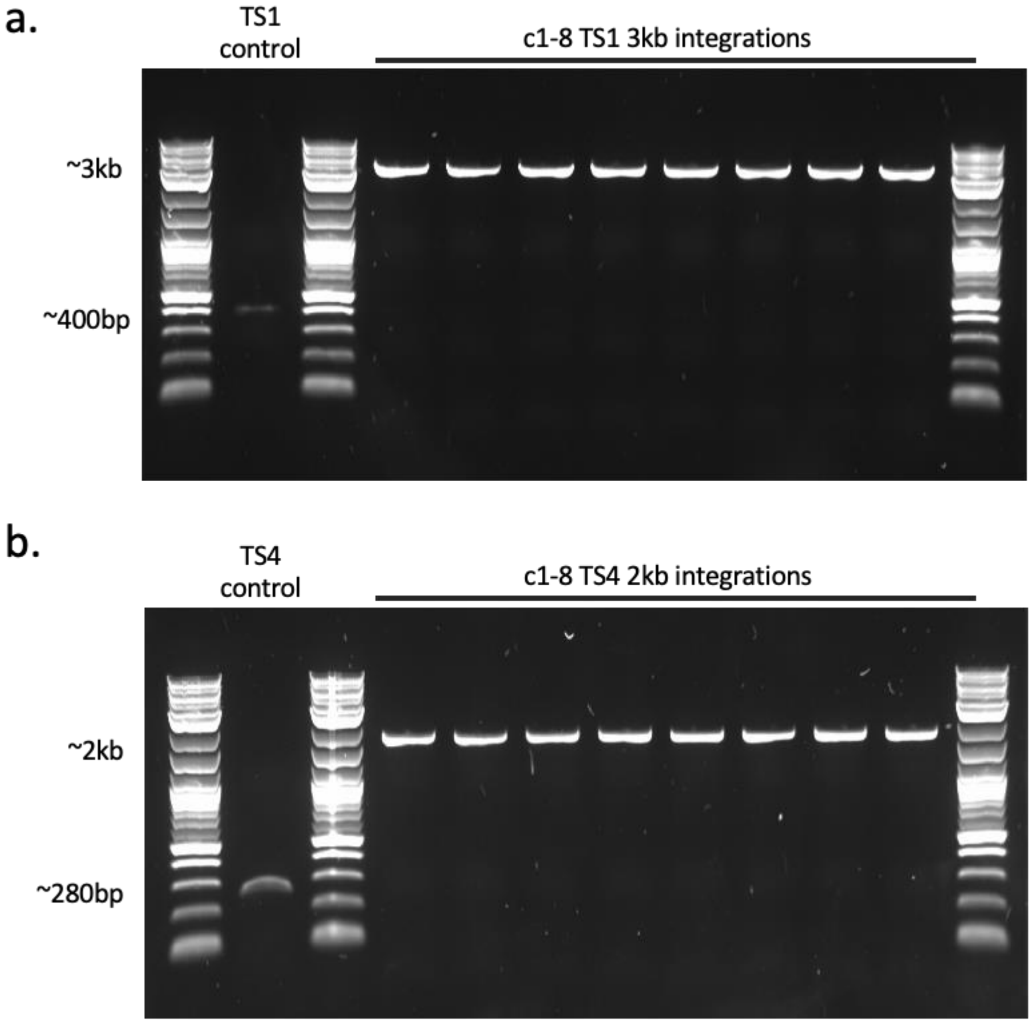


Figure S14. Genotypes of iterative integrations with pTarget-TS1 and pTarget-TS4. Eight colonies were screened in the TS1 and TS4 sites showing integrations of both a 3-kb cargo in TS1 and 2-kb cargo in TS4.

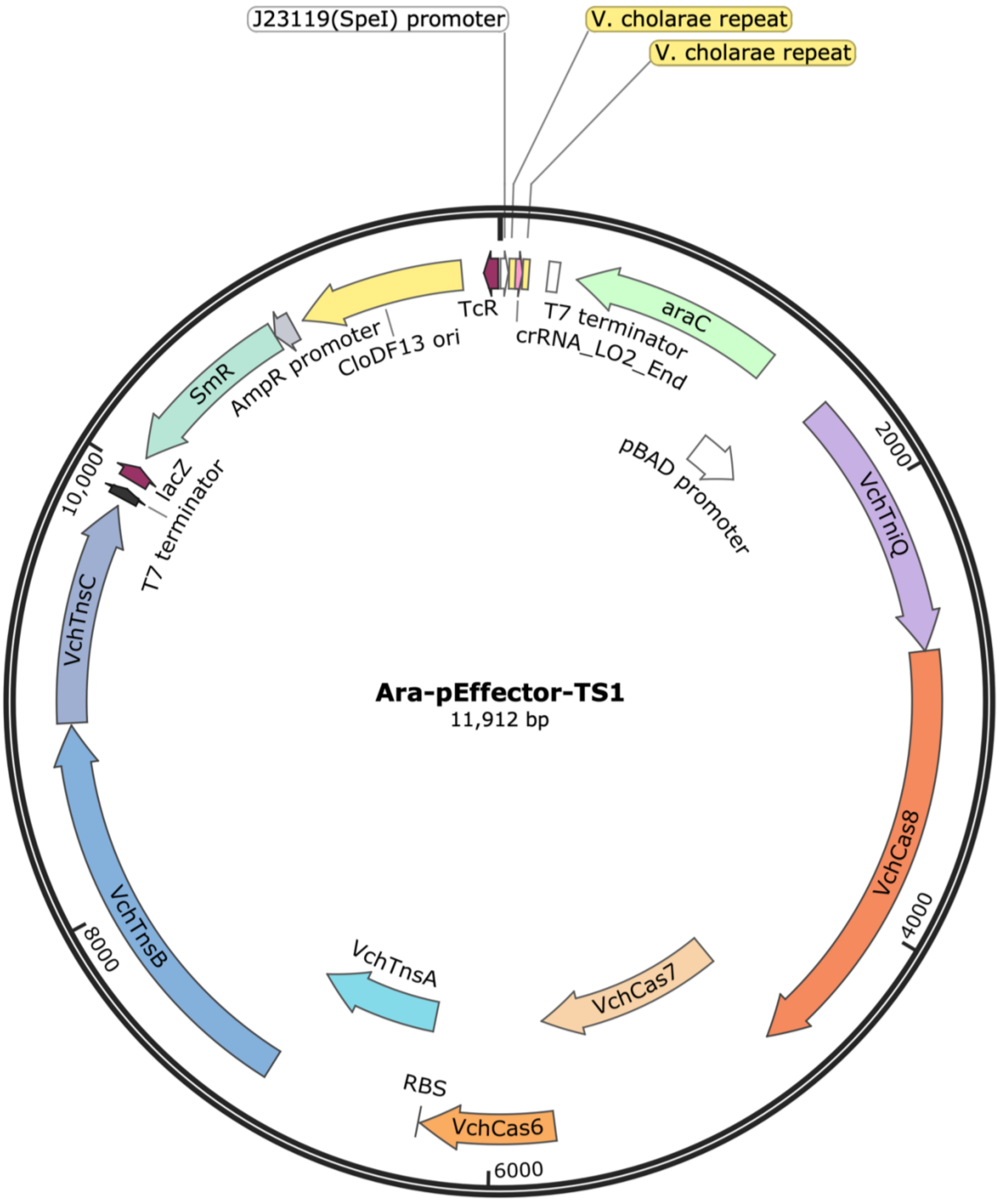


Figure S15. Representative plasmid map of Ara-pEffector-TS1. Sequence data for this plasmid and the rest can be found at https://github.com/meganwang08/A-universal-system-for-streamlined-genome-integrations-with-CRISPR-associated-transposases.

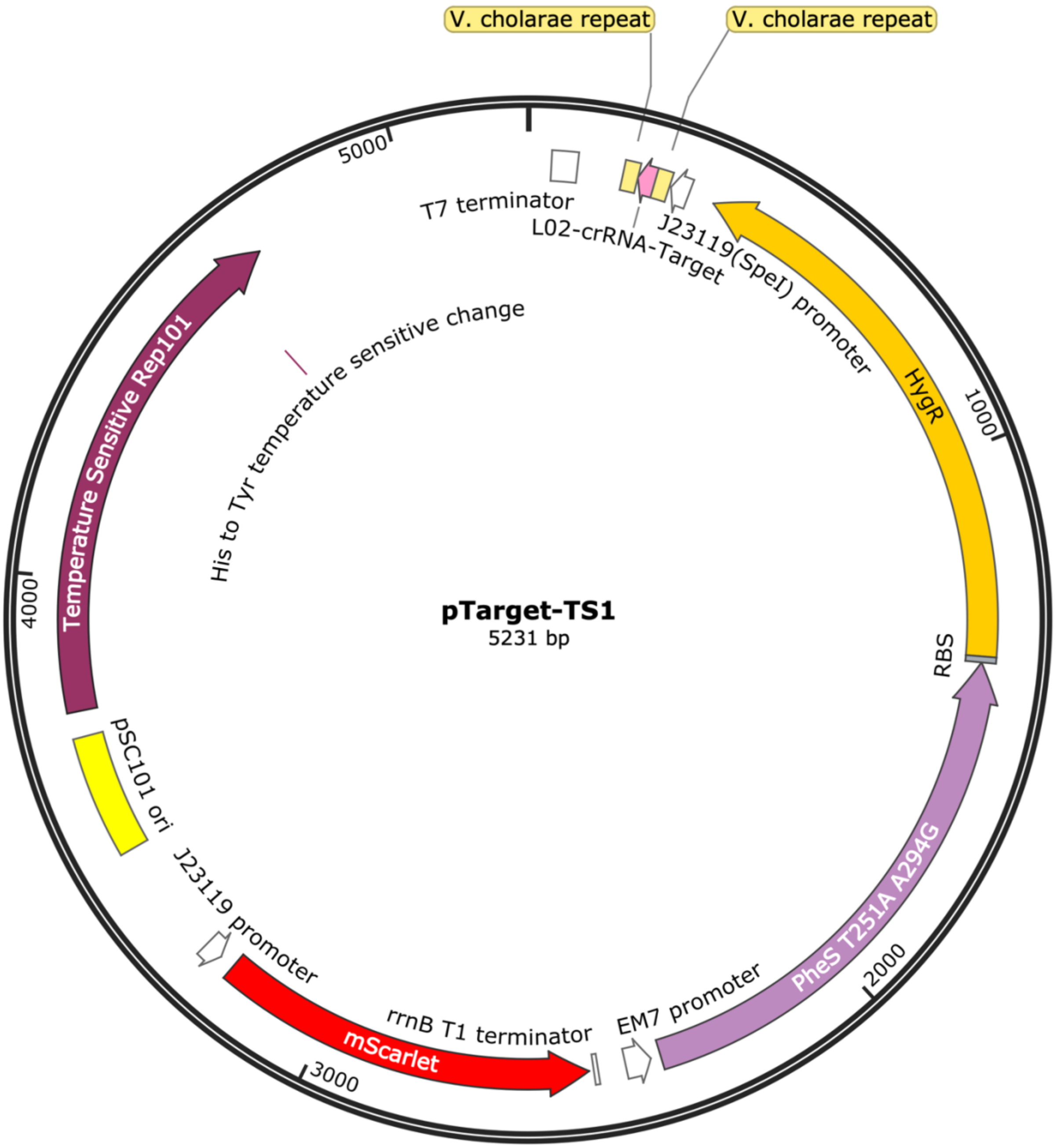


Figure S16. Representative plasmid map of pTarget-TS1. Sequence data for this plasmid and the rest can be found at https://github.com/meganwang08/A-universal-system-for-streamlined-genome-integrations-with-CRISPR-associated-transposases.

Figure S17. Annotated sequence of Ara-pEffector-TS1. J23119 promoter, V. cholarae repeat, TS1-crRNA-target, araC-pBad promoter, and Vch-CAST comprised of (TniQ, Cas5-8, Cas7, Cas6, TnsA, TnsB, TnsC)

ttgacagctagctcagtcctaggtataatactagtCAcGTGAACTGCCGAGTAGGTAGCTGATAACaccagacccgcgagca
ttaattcttgcctccaGTGAACTGCCGAGTAGGTAGCTGATAACtcgagagatccggctgctaacaaacccgcttaaggaagat
ggccaacttatttacataaggctgctgccaccgctgagcaataactagcataaccccttggggcctctaaacgggtcttgaggggttttttggtttaaaccca
tctaattgggaattcccccctgcctcgcgcgtttcggtgatgacggtgaaaacctctgacacaGCGATCGCTTATGACAACTTGACG
GCTACATCATTCACTTTTTCTTCACAACCGGCACGGAACTCGCTCGGGCTGGCCCCGGTGCAT
TTTTTAAATACCCGCGAGAAATAGAGTTGATCGTCAAAACCAACATTGCGACCGACGGTGGC
GATAGGCATCCGGGTGGTGCTCAAAAGCAGCTTCGCCTGGCTGATACGTTGGTCCTCGCGCC
AGCTTAAGACGCTAATCCCTAACTGCTGGCGGAAAAGATGTGACAGACGCGACGGCGACAA
GCAAACATGCTGTGCGACGCTGGCGATATCAAAATTGCTGTCTGCCAGGTGATCGCTGATGT
ACTGACAAGCCTCGCGTACCCGATTATCCATCGGTGGATGGAGCGACTCGTTAATCGCTTCC
ATGCGCCGCAGTAACAATTGCTCAAGCAGATTTATCGCCAGCAGCTCCGAATAGCGCCCTTC
CCCTTGCCCGGCGTTAATGATTTGCCCAAACAGGTCGCTGAAATGCGGCTGGTGCGCTTCAT
CCGGGCGAAAGAACCCCGTATTGGCAAATATTGACGGCCAGTTAAGCCATTCATGCCAGTA
GGCGCGCGGACGAAAGTAAACCCACTGGTGATACCATTCGCGAGCCTCCGGATGACGACCG
TAGTGATGAATCTCTCCTGGCGGGAACAGCAAAATATCACCCGGTCGGCAAACAAATTCTCG
TCCCTGATTTTTCACCACCCCCTGACCGCGAATGGTGAGATTGAGAATATAACCTTTCATTCC
CAGCGGTCGGTCGATAAAAAAATCGAGATAACCGTTGGCCTCAATCGGCGTTAAACCCGCC
ACCAGATGGGCATTAAACGAGTATCCCGGCAGCAGGGGATCATTTTGCGCTTCAGCCATACT
TTTCATACTCCCGCCATTCAGAGAAGAAACCAATTGTCCATATTGCATCAGACATTGCCGTC
ACTGCGTCTTTTACTGGCTCTTCTCGCTAACCAAACCGGTAACCCCGCTTATTAAAAGCATTC
TGTAACAAAGCGGGACCAAAGCCATGACAAAAACGCGTAACAAAAGTGTCTATAATCACGG
CAGAAAAGTCCACATTGATTATTTGCACGGCGTCACACTTTGCTATGCCATAGCATTTTTATC

CATAAGATTAGCGGATCCTACCTGACGCTTTTTATCGCAACTCTCTACTGTTTCTCCATACCC

GGGATCCGAATTCGAGCGAAGGAGATATACATATGTTTTTGCAAAGACCTAAACCTTACAGC

GATGAAAGtTTAGAAAGtTTCTTTATCCGAGTGGCTAACAAAAATGGCTACGGTGATGTCCAT

CGCTTCCTAGAAGCCACTAAACGATTCCTTCAAGACATTGACCATAATGGCTATCAAACCTT

TCCGACTGATATAACTCGGATAAACCCATACTCAGCTAAAAACAGTTCCAGCGCACGAACTG

CGTCATTCCTGAAaCTTGCACAATTGACATTTAATGAACCGCCAGAGCTACTTGGGTTGGCAA

TTAACAGAACAAACATGAAATACTCGCCGTCAACTAGCGCGGTTGTTAGAGGTGCAGAAGT

CTTTCCTCGCAGTTTACTACGGACGCACTCCATCCCCTGCTGTCCTTTGTGTCTGCGAGAAAA

TGGCTACGCCTCCTACCTTTGGCACTTTCAGGGGTACGAATACTGCCACAGCCATAACGTAC

CTTTAATTACCACTTGTAGCTGTGGTAAGGAGTTTGACTACCGAGTATCTGGGTTAAAGGGC

ATTTGCTGCAAATGCAAGGAGCCTATCACCTTAACCAGCAGGGAGAACGGTCATGAGGCAG

CGTGTACTGTTTCAAACTGGCTTGCTGGCCATGAATCTAAACCTCTGCCAAATCTTCCTAAAA

GCTACCGATGGGGTTTAGTTCATTGGTGGATGGGTATTAAAGATAGCGAgTTCGATCACTTTT

CGTTCGTTCAATTTTTCTCAAACTGGCCAAGGTCATTCCACTCGATAATCGAAGATGAAGTA

GAGTTCAACCTTGAGCATGCTGTTGTCAGCACGTCTGAATTACGACTAAAAGATCTTCTTGG

TCGATTGTTTTTCGGTTCAATTCGGTTACCTGAGCGGAATCTTCAACACAATATCATCCTTGG

TGAGCTTCTCTGCTATTTAGAAAATCGTTTATGGCAAGACAAGGGATTAATCGCCAACCTCA

AAATGAACGCGTTAGAGGCGACTGTAATGTTAAATTGTAGCCTCGATCAGATTGCATCAATG

GTTGAACAACGCATCTTGAAGCCAAATCGAAAAAGCAAGCCCAACAGCCCTCTTGATGTTAC

CGATTATCTATTTCATTTCGGCGATATTTTCTGTCTTTGGTTAGCTGAgTTCCAAAGCGATGAG

TTTAACCGTTCGTTTTATGTGTCGAGGTGGTAAATGCAAACTCTGAAAGAACTAATCGCATC

CAATCCTGACGACTTAACAACTGAACTTAAGAGAGCATTTCGTCCACTCACACCGCATATTG

CAATTGATGGTAATGAACTTGACGCACTGACGATATTAGTCAATTTAACCGATAAGACTGAT

GATCAGAAAGACCTGCTCGATCGAGCCAAATGCAAGCAAAAACTTCGAGATGAAAAATGGT

GGGCTAGCTGCATAAATTGCGTTAAtTACAGACAAAGCCATAACCCAAAATTCCCGGATATA

CGTTCTGAAGGCGTGATTCGAACCCAAGCCCTgGGTGAATTACCGAGTTTCCTACTCTCGTCc

TCAAAAATCCCACCATACCATTGGTCATATAGCCATGATTCAAAATACGTCAACAAAAGCGC
ATTCCTCACCAATGAGTTTTGTTGGGATGGTGAGATCTCATGTTTGGGTGAGCTTCTTAAAGA
TGCAGATCACCCACTTTGGAATACTTTGAAAAAGTTAGGTTGTTCTCAAAAAACTTGCAAAG
CAATGGCAAAACAACTAGCTGATATTACTCTCACGACTATCAATGTCACACTCGCACCAAAT
TACCTGACTCAAATATCTCTCCCCGATAGTGATACATCCTACATTTCACTCTCTCCTGTTGCA
TCGCTATCGATGCAAAGCCACTTTCATCAGAGGCTCCAAGATGAAAATCGGCATAGCGCGAT
AACGCGGTTTAGCCGAACTACCAACATGGGGGTCACTGCGATGACATGTGGCGGCGCATTTA
GAATGTTAAAGTCTGGCGCTAAGTTTTCCAGCCCCCCTCATCACCGATTAAACAGTAAACGA
AGTTGGTTgACGTCAGAGCATGTTCAGTCATTAAAACAGTACCAGCGCCTCAACAAAAGCCT
CATACCTGAAAACTCTCGGATTGCACTCCGTAGAAAATACAAAATCGAGCTTCAAAATATGG
TCAGATCTTGGTTTGCAATGCAAGACCATACCCTTGATTCGAACATACTTATCCAACATTTGA
ATCATGACCTATCTTACTTAGGAGCCACAAAACGTTTTGCATACGATCCAGCGATGACCAAG
CTCTTTACTGAGCTTTTGAAACGAGAGTTATCAAATTCAATCAATAATGGTGAGCAACACAC
TAATGGATCGTTTTTAGTCCTACCGAATATCAGGGTTTGTGGCGCAACAGCTTTAAGCTCCCC
GGTAACGGTGGGGATtCCATCACTTACAGCTTTCTTTGGCTTCGTTCACGCATTTGAACGGAA
TATAAATCGCACCACCTCATCGTTTCGTGTTGAATCCTTTGCGATATGCGTCCATCAACTACA
TGTCGAGAAGCGAGGTTTGACAGCAGAGTTTGTGGAAAAAGGCGACGGGACTATATCCGCT
CCCGCGACCCGGGATGACTGGCAGTGTGATGTCGTATTTAGCCTTATTTTGAACACCAACTTT
GCTCAACATATTGACCAAGATACGTTAGTTACATCACTACCAAAGCGATTGGCTCGgGGTTC
AGCAAAAATTGCGATTGATGACTTTAAACATATCAACTCATTCTCGACATTAGAAACAGCGA
TCGAATCTCTGCCAATAGAAGCTGGTAGGTGGTTATCACTTTACGCACAGTCAAACAATAAT
CTAAGTGATCTATTAGCAGCCATGACAGAGGACCATCAGCTCATGGCAAGCTGCGTCGGTTA
CCACTTGTTAGAAGAGCCCAAAGATAAACCAAACTCCCTCAGAGGTTACAAACACGCTATC
GCCGAGTGCATCATTGGACTCATTAACTCAATCACCTTTAGCTCAGAGACTGATCCCAACAC
AATCTTTTGGTCGCTAAAGAACTATCAAAACTACCTAGTGGTACAGCCAAGGAGTATCAACG
ATGAAACTACCGACAAATCTAGCCTATGAGCGCTCTATCGACCCATCAGATGTCTGTTTTTTT

GTCGTCTGGCCCGATGATAGAAAAACACCTTTAACCTACAATTCTCGTACTCTGCTCGGGCA

AATGGAAGCGGCATCATTAGCCTATGATGTCTCAGGTCAACCAATAAAAAGTGCCACCGCTG

AGGCGTTAGCTCAAGGGAACCCTCATCAAGTTGATTTCTGCCACGTTCCATAcGGTGCGAGT

CATATTGAATGCAGTTTCTCCGTCTCGTTTTCTTCTGAACTACGTCAACCATATAAGTGTAAC

TCAAGCAAAGTTAAACAAACGCTAGTGCAATTAGTCGAGCTCTACGAAACGAAAATCGGCT

GGACTGAGCTAGCAACCCGATATTTGATGAAcATTTGCAACGGTAAATGGCTGTGGAAAAAT

ACCCGTAAAGCcTATTGCTGGAACATTGTACTTACACCTTGGCCtTGGAACGGGGAAAAGGTT

GGATTTGAAGATATtCGTACTAACTACACCTCACGGCAAGACTTTAAAAATAATAAAAATTG

GTCTGCTATAGTTGAAATGATCAAAACCGCATTTTCTAGTACTGATGGGCTGGCGATATTTG

AAGTCAGGGCCACCTTGCACTTGCCAACGAATGCTATGGTGCGGCCAAGCCAAGTTTTCACA

GAAAAAGAAAGTGGCAGTAAAAGTAAATCTAAAACTCAAAACAGTCGAGTTTTTCAGAGTA

CAACTATTGATGGTGAACGATCGCCAATACTAGGGGCCTTTAAAACGGGAGCAGCTATTGCA

ACCATTGACGACTGGTATCCTGAAGCCACTGAGCCACTAAGGGTCGGACGGTTTGGGGTTCA

TCGCGAAGATGTCACTTGCTACCGTCATCCGTCTACCGGAAAAGATTTTTTCTCGATATTACA

ACAAGCAGAGCACTATATTGAAGTGTTGAGCGCCAACAAAACTCCCGCTCAAGAAACTATC

AACGACATGCACTTTTTAATGGCTAACCTGATTAAGGGTGGGATGTTCCAGCATAAAGGAGA

CTGACTGTGAAATGGTATTATAAGACAATCACCTTTCTGCCAGAGTTGTGCAACAACGAGTC

ACTGGCTGCAAAGTGTCTcCGCGTTCTGCATGGATTTAACTATCAGTATGAGACACGAAATA

TcGGCGTTTCATTTCCGCTTTGGTGTGATGCAACGGTTGGAAAAAAGATTTCATTTGTCAGCA

AGAACAAGATAGAACTCGACTTACTACTTAAACAACACTATTTCGTCCAAATGGAACAACTT

CAATAcTTTCATATATCCAACACTGTTCTCGTCCCAGAAGATTGTACATACGTTTCCTTTAGA

CGCTGTCAATCTATAGATAAGCTCACAGCAGCAGGGCTGGCAAGGAAAATCAGACGCCTGG

AGAAACGTGCTCTATCTAGAGGCGAGCAATTTGACCCATCATCTTTTGCTCAAAAAGAGCAT

ACTGCAATAGCGCACTACCACTCACTTGGGGAGTCCAGCAAACAGACGAACCGCAACTTTC

GACTCAATATCAGGATGCTCTCGGAGCAACCCCGTGAGGGGAACTCGATTTTTAGTAGCTAT

GGCTTATCAAATTCAGAAAACTCGTTTCAGCCTGTACCCTTAATCTGACCTTTAATAAGGAG

ATATACCATGGcgACAAGtTTACCTACGCCCTCAGCAATTACGACTTCGGCGTTAGAGTATGC

ATTCCATACTCCCGCTCGCAATCTAACGAAATCTCGCGGAAAAAAcATTCATCGTTATGTCAG

TGTAAAGATGAGTAAGAGGATTACGGTAGAATCTACTCTAGAGTGTGATGCCTGCTATCACT

TTGATTTTGAGCCAAGTATTGTTCGCTTTTGCGCTCAACCGATTCGATTTTTATATTATCTCAA

TGGTCAGTCTCACTCCTATGTTCCTGACTTTCTAGTTCAATTTGATACCAACGAGTTTGTTCTA

TATGAAGTAAAGTCAGCTTATGCTAAGAACAAACCTGATTTTGATGTTGAATGGGAGGCGAA

AGTAAAAGCAGCAACTGAACTAGGGCTAGAATTGGAGCTTGTTGAAGAGAGTGATATTAGG

GATACGGTTGTATTAAATAATCTTAAGCGCATGCATCGTTATGCTTCGAAAGATGAGCTGAA

TAACGTACATAACTCTCTCTTAAAAATAATAAAGTACAATGGCGCCCAATCTGCAAGATGCT

TGGGAGAACAGTTGGGTTTAAAAGGCCGAACTGTTTTACCAATTTTGTGCGATTTGCTGTCA

AGGTGTTTACTCGATACACGTTTGGATAAGCCTCTATCTCTTGAATCTCGATTTGAGTTGGCC

AGTTATGGCTAAGAAAGGGTTCTCaAGTTTCCATAGAAAAGCAGTCTCTTCCCAAGATACGC

TTGAATCGATAGAGCTTGTCTCTAGCGCTAATTGTCTAGAAAGTGTTACGTATCAAGATATA

TCAGCATTTCCCGAAACAATTGCGGTAGAGATTAATTTCCGATTAAGCATTCTTCGCTTTTTA

GCGCGAAAGTGCGAAACCATTGTGGCCAAATCAATTGAACCACATCGTGTAGAGCTACAGC

AAAACTATAGTAGAAAAATACCCAGTGCAATAACGATATATCGATGGTGGCTTGCTTTTCGA

AAATCAGACTACAACCCCATTAGCTTAGCACCTAATATCAAGGATAGAGGTAATAGAGAGA

CAAAAGTGTCAACAGTTGTTGATTCTATTATGGAACAGGCAGTTGAAAGAGTTATATCTGGA

CGAAAAGTCAATGTTAGCTCTGCATATAAACGTGTTCGACGAAAAGTTCGTCAATACAATCT

AACTCATGGAACGAAATACACGTATCCTAAGTACGAATCTGTAAGAAAGCGAGTAAAAAAG

AAAACCCCATTTGAGTTATTAGCCGCAGGGAAAGGGGAGAGAGTAGCTAAGAGAGAGTTTC

GCCGAATGGGAAAAAAGATCCTCACGTCTAGCGTGCTAGAGAGGGTTGAAATAGATCACAC

TGTCGTTGACCTTTTTGCAGTACACGAAGAGTATCGAATcCCATTGGGCCGACCTTGGCTTAC

TCAATTGGTTGATTGTTACAGTAAAGCTGTAATCGGTTTTTATTTAGGTTTCGAGCCTCCTAG

CTATGTGTCGGTTTCCCTTGCACTTAAGAATGCAATACAACGCAAAGATGACTTAATCTCCTC

GTATGAATCGATCGAGAATGAATGGCTATGTTATGGcATCCCAGACCTACTCGTAACTGATA

ATGGTAAAGAGTTTTTGTCGAAAGCATTTGATCAAGCATGTGAATCACTATTGATCAATGTG

CATCAAAATAAAGTTGAGACGCCCGACAACAAACCTCATGTTGAACGTAACTACGGGACTA

TTAATACTTCTCTGTTAGACGATTTACCTGGGAAATCCTTCAGCCAGTACCTTCAAAGAGAA

GGGTACGACTCTGTGGGAGAAGCTACCCTTACACTCAATGAGATTAGAGAAATTTACTTAAT

TTGGTTGGTGGATATTTATCATAAAAAACCCAATCAGAGAGGCACTAATTGTCCTAATGTTG

CGTGGAAAAAGGGTTGTCAAGAATGGGAACCAGAGGAGTTCTCTGGTTCTAAAGACGAATT

AGACTTTAAATTTGCTATTGTTGATTACAAACAACTTACTAAAGTAGGGATAACTGTCTACA

AAGAACTGAGTTATAGCAATGACCGTTTAGCTGAATATAGAGGGAAGAAAGGAAACCATAA

AGTTCAGTTCAAGTATAACCCTGAGTGTATGGCAGTTATTTGGGTGTTGGATGAGGATATGA

ATGAGTACTTTACAGTTAATGCGATTGACTACGAATATGCAAGTAGAGTATCACTTTGGCAA

CATAAATATAACATGAAATATCAAGCAGAACTAAATTCAGCAGAATATGATGAgGACAAGG

AAATTGATGCAGAAATAAAAATTGAAGAAATCGCAGATCGTTCAATTGTTAAGACTAACAA

AATCAGAGCTCGGAGGCGTGGCGCTAGGCATCAAGAGAATAGCGCAAGGGCTAAGTCAATC

AGTAATGCGAACCCGGCCTCGATACAAAAACATGAAGATGAAATCGTTAGTGCAGATAATG

ACGATTGGGATATTGATTATGTCTGAGAAATCGTCAAATGAGTGAAACGCGTGAGGCTCGTA

TATCAAGAGCTAAAAGGGCATTTGTATCCACACCGAGCGTTAGGAAAATCTTGAGTTACATG

GATAGATGTAGAGAtCTATCAGACCTAGAGTCTGAGCCTACATGCATGATGGTCTATGGTGC

TTCGGGTGTAGGTAAAACGACCGTCATCAAGAAATACTTAAATCAAAACAGAAGAGAGTCC

GAAGCCGGGGGCGATATAATACCGGTTTTGCATATTGAGTTGCCAGACAATGCGAAGCCAG

TAGATGCAGCAAGGGAATTGCTGGTTGAAATGGGTGACCCGCTAGCACTTTATGAAACTGAC

TTAGCTAGATTGACGAAAAGACTGACTGAATTAATCCCTGCGGTCGGCGTGAAGCTGATTAT

TATCGATGAGTTCCAACATTTGGTGGAAGAAAGGTCAAATCGGGTTCTTACCCAAGTAGGTA

ATTGGCTAAAAATGATACTTAACAAAACGAAATGTCCAATTGTTATATTTGGTATGCCATAC

TCAAAAGTTGTACTGCAAGCAAACTCGCAACTTCACGGGCGATTTTCCATTCAGGTTGAACT

GCGCCCCTTTAGCTACCAGGGAGGTAGAGGTGTATTTAAAACTTTTTTGGAATACCTTGATA

AAGCCCTACCTTTTGAAAAACAGGCTGGCTTAGCCAACGAAAGTTTGCAGAAAAAATTGTAT

GCATTCTCTCAGGGAAACATGCGTTCGTTGAGAAACCTTATTTATCAAGCATCTATCGAAGC

AATTGATAATCAGCATGAGACGATAACCGAAGAAGATTTCGTTTTTGCATCGAAGTTGACAT

CGGGCGATAAACCCAACTCATGGAAAAATCCTTTTGAGGAGGGTGTTGAGGTAACAGAAGA

TATGTTACGACCGCCACCAAAAGATATTGGTTGGGAgGACTATTTGAGACATTCAACCCCGA

GAGTGAGTAAACCAGGTAGAAATAAAAACTTTTTCGAATAACCTAGGCTGCTGCGCTGCTGC

CACCGCTGAGCAATAACTAGCATAACCCCTTGGGGCCTCTAAACGGGTCTTGAGGGGTTTTT

TGCTGAAACCTCAGGCATTgaacctgcacGGTACCGAGCTCGAATTCACTGGCCGTCGTTTTACAA

CGTCGTGACTGGGAAAACCCTGGCGTTACCCAACTTAATCGCCTTGCtATCGTGGCCGGATCT

TGCGGCCCCTCGGCTTGAACGAATTGTTAGACATTATTTGCCGACTACCTTGGTGATCTCGCC

TTTCACGTAGTGGACAAATTCTTCCAACTGATCTGCGCGCGAGGCCAAGCGATCTTCTTCTTG

TCCAAGATAAGCCTGTCTAGCTTCAAGTATGACGGGCTGATACTGGGCCGGCAGGCGCTCCA

TTGCCCAGTCGGCAGCGACATCCTTCGGCGCGATTTTGCCGGTTACTGCGCTGTACCAAATG

CGGGACAACGTAAGCACTACATTTCGCTCATCGCCAGCCCAGTCGGGCGGCGAGTTCCATAG

CGTTAAGGTTTCATTTAGCGCCTCAAATAGATCCTGTTCAGGAACCGGATCAAAGAGTTCCT

CCGCCGCTGGACCTACCAAGGCAACGCTATGTTCTCTTGCTTTTGTCAGCAAGATAGCCAGA

TCAATGTCGATCGTGGCTGGCTCGAAGATACCTGCAAGAATGTCATTGCGCTGCCATTCTCC

AAATTGCAGTTCGCGCTTAGCTGGATAACGCCACGGAATGATGTCGTCGTGCACAACAATGG

TGACTTCTACAGCGCGGAGAATCTCGCTCTCTCCAGGGGAAGCCGAAGTTTCCAAAAGGTCG

TTGATCAAAGCTCGCCGCGTTGTTTCATCAAGCCTTACGGTCACCGTAACCAGCAAATCAAT

ATCACTGTGTGGCTTCAGGCCGCCATCCACTGCGGAGCCGTACAAATGTACGGCCAGCAACG

TCGGTTCGAGATGGCGCTCGATGACGCCAACTACCTCTGATAGTTGAGTCGATACTTCGGCG

ATCACCGCTTCCCTCATACTCTTCCTTTTTCAATATTATTGAAGCATTTATCAGGGTTATTGTC

TCATGAGCGGATACATATTTGAATGTATTTAGAAAAATAAACAAATAGCTAGCTCACTCGGT

CGCTACGCTCCGGGCGTGAGACTGCGGCGGGCGCTGCGGACACATACAAAGTTACCCACAG

ATTCCGTGGATAAGCAGGGGACTAACATGTGAGGCAAAACAGCAGGGCCGCGCCGGTGGCG

TTTTTCCATAGGCTCCGCCCTCCTGCCAGAGTTCACATAAACAGACGCTTTTCCGGTGCATCT

GTGGGAGCCGTGAGGCTCAACCATGAATCTGACAGTACGGGCGAAACCCGACAGGACTTAA
AGATCCCCACCGTTTCCGGCGGGTCGCTCCCTCTTGCGCTCTCCTGTTCCGACCCTGCCGTTT
ACCGGATACCTGTTCCGCCTTTCTCCCTTACGGGAAGTGTGGCGCTTTCTCATAGCTCACACA
CTGGTATCTCGGCTCGGTGTAGGTCGTTCGCTCCAAGCTGGGCTGTAAGCAAGAACTCCCCG
TTCAGCCCGACTGCTGCGCCTTATCCGGTAACTGTTCACTTGAGTCCAACCCGGAAAAGCAC
GGTAAAACGCCACTGGCAGCAGCCATTGGTAACTGGGAGTTCGCAGAGGATTTGTTTAGCTA
AACACGCGGTTGCTCTTGAAGTGTGCGCCAAAGTCCGGCTACACTGGAAGGACAGATTTGGT
TGCTGTGCTCTGCGAAAGCCAGTTACCACGGTTAAGCAGTTCCCCAACTGACTTAACCTTCG
ATCAAACCACCTCCCCAGGTGGTTTTTTCGTTTACAGGGCAAAAGATTACGCGCAGAAAAAA
AGGATCTCAAGAAGATCCTTTGATCTTTTCTACTGAACCGCTCTAGATTTCAGTGCAATTTAT
CTCTTCAAATGTAGCACCTGAAGTCAGCCCCATACGATATAAGTTGTAATTCTCATGTTAGTC
ATGCCCCGCGCCCACCGGAAGGAGCTGACTGGGTTGAAGGCTCTCAAGGGCATCGGTCGcacg
cttgtg

Figure S18. Annotated sequence of pTarget -TS1. J23119 promoter, V. cholarae repeat, TS1-crRNA-target, mScarlet, HygroR-PheS

ttgacagctagctcagtcctaggtataatactagtCAcGTGAACTGCCGAGTAGGTAGCTGATAACaccagacccgcgagca
ttaattcttgcctccaGTGAACTGCCGAGTAGGTAGCTGATAACtcgagagatccggctgctaacaaacccgcttaaggaagat
ggccaacttatttacataaggctgctgccaccgctgagcaataactagcataaccccttggggcctctaaacgggtcttgaggggttttttggtttaaaccca
tctaattgggaattcccctgcctcgGTTAACtcactgtacactacgccttttgtggagatgtctaataaactaaaaatttttactttccgctgcataaccc
tgcttcggggtcattatagcgatttttcggtatatccatcctttttcgcacgatatacaggattttgccaaagggttcgtgtagactttccttggtgtatccaacg
gcgtcagccgggcaggataggtgaagtaggcccacccgcgagcgggtgttccttcttcactgtcccttattcgcacctggcggtgctcaacgggaatcct
gctctgcgaggctggccgataagctctgataacgcaggaaagaacatgggcagttcaacctgttgataCTCGAGggcacatagccttgctcaaatt
ggaatcaggtttgtgccaataccagtagaaacagacgaagaatccatgggtatggacagttttccctttgatatgtaacggtgaacagttgttctacttttgttt
gttagtcttgatgcttcactgatagatacaagagccataagaacctcagatccttccgtatttagccagtatgttctctagtgtggttcgttgttttttgcgtgagcc
atgagaacgaaccattgagatcatacttactttgcatgtcactcaaaaattttgcctcaaaactggtgagctgaatttttgcagttaaagcatcgtgtagtgttttt
cttagtccgttacgtaggtaggaatctgatgtaatggttgttggtattttgtcaccattcatttttatctggttgttctcaagttcggttacgagatccatttgtctatc
tagttcaacttggaaaatcaacgtatcagtcgggcggcctcgcttatcaaccaccaatttcatattgctgtaagtgtttaaatctttacttattggtttcaaaaccc
attggttaagccttttaaactcatggtagttattttcaagcattaacatgaacttaaattcatcaaggctaatctctatatttgccttgtgagttttcttttgtgttagttc
ttttaataaccactcataaatcctcatagagtatttgttttcaaaagacttaacatgttccagattatattttatgaatttttttaactggaaaagataaggcaatatct
cttcactaaaaactaattctaatttttcgcttgagaacttggcatagtttgtccactggaaaatctcaaagcctttaaccaaaggattcctgatttccacagttctc
gtcatcagctctctggttgctttagctaatacaccataagcattttccctactgatgttcatcatctgagcgtattggttataagtgaacgataccgtccgttctttc
cttgtagggttttcaatcgtggggttgagtagtgccacacagcataaaattagcttggtttcatgctccgttaagtcatagcgactaatcgctagttcatttgctt
tgaaaacaactaattcagacatacatctcaattggtctaggtgattttaatcactataccaattgagatgggctagtcaatgataattactagtccttttcctttga
gttgtgggtatctgtaaattctgctagacctttgctggaaaacttgtaaattctgctagaccctctgtaaattccgctagacctttgtgtgttttttttgtttatattca
agtggttataatttatagaataaagaaagaataaaaaaagataaaaagaatagatcccagccctgtgtataactcactactttagtcagttccgcagtattaca
aaaggatgtcgcaaacgctgtttgctcctctacaaaacagaccttaaaaccctaaaggcttaagtagcaccctcgcaagctcggttgcggccgcaatcgg
gcaaatcgctgaatattccttttgtctccgaccatcaggcacctgagtcgctgtctttttcgtgacattcagttcgctgcgctcacTCGCGATTTACA
GCTAGCTCAGTCCTAGGTATAATGCTAGCTCTAGAGATTAAAGAGGAGAAATACTAGATGGT

CAGTAAAGGCGAAGCAGTTATCAAAGAGTTCATGCGCTTCAAAGTTCATATGGAAGGGTCG

ATGAACGGGCACGAATTTGAAATTGAAGGCGAAGGCGAAGGCCGCCCATATGAAGGGACCC

AAACCGCAAAGCTTAAGGTTACTAAAGGCGGTCCATTACCCTTTTCGTGGGACATTTTAAGC

CCACAGTTTATGTACGGGAGTCGCGCTTTCATCAAGCACCCTGCGGACATCCCAGATTACTA

CAAACAGTCTTTCCCCGAGGGGTTCAAGTGGGAGCGCGTGATGAACTTCGAGGATGGCGGA

GCCGTGACGGTCACCCAAGATACCTCTTTGGAGGACGGTACGTTGATCTACAAAGTGAAATT

GCGTGGCACGAATTTTCCACCTGATGGGCCTGTCATGCAGAAAAAGACAATGGGATGGGAA

GCTTCCACGGAGCGCCTTTACCCAGAGGACGGTGTTCTTAAAGGGGATATCAAAATGGCGCT

GCGTCTTAAAGATGGAGGCCGCTACCTGGCGGACTTCAAGACTACTTACAAGGCCAAAAAA

CCAGTGCAGATGCCGGGTGCGTACAATGTAGATCGTAAATTAGATATTACAAGTCACAATGA

AGATTACACGGTCGTAGAGCAGTATGAGCGCAGTGAGGGGCGTCACTCTACGGGCGGTATG

GACGAGTTATACAAGTAAGCGGCCGCcaaataaaGAGCTCcgcaaaaaccccgcttcggcggggtttttcgcACGC

GTcacgtGTTGACAATTAATCATCGGCATAGTATATCGGCATAGTATAATACGACAAGGTGAGG

AACTAAACCATGTCACATCTCGCAGAACTGGTTGCCAGTGCGAAGGCGGCCATTAGCCAGG

CGTCAGATGTTGCCGCGTTAGATAATGTGCGCGTCGAATATTTGGGTAAAAAAGGGCACTTA

ACCCTTCAGATGACGACCCTGCGTGAGCTGCCGCCAGAAGAGCGTCCGGCAGCTGGTGCGG

TTATCAACGAAGCGAAAGAGCAGGTTCAGCAGGCGCTGAATGCGCGTAAAGCGGAACTGGA

AAGCGCTGCACTGAATGCGCGTCTGGCGGCGGAAACGATTGATGTCTCTCTGCCAGGTCGTC

GCATTGAAAACGGCGGTCTGCATCCGGTTACCCGTACCATCGACCGTATCGAAAGTTTCTTC

GGTGAGCTTGGCTTTACCGTGGCAACCGGGCCGGAAATCGAAGACGATTATCATAACTTCGA

TGCTCTGAACATTCCTGGTCACCACCCGGCGCGCGCTGACCACGACACTTTCTGGTTTGACA

CTACCCGCCTGCTGCGTACCCAGACCTCTGGCGTACAGATCCGCACCATGAAAGCCCAGCAG

CCACCGATTCGTATCATCGCGCCTGGCCGTGTTTATCGTAACGACTACGACCAGACTCACAC

GCCGATGTTCCATCAGATGGAAGGTCTGATTGTTGATACCAACATCAGCTTTACCAACCTGA

AAGGCACGCTGCACGACTTCCTGCGTAACTTCTTTGAGGAAGATTTGCAGATTCGCTTCCGT

CCTTCCTACTTCCCGTTTGCCGAACCTTCTGCAGAAGTGGACGTCATGGGTAAAAACGGTAA

ATGGCTGGAAGTGCTGGGCTGCGGGATGGTGCATCCGAACGTGTTGCGTAACGTTGGCATCG
ACCCGGAAGTTTACTCTGGTTTCGGCTTCGGGATGGGGATGGAGCGTCTGACTATGTTGCGT
TACGGCGTCACCGACCTGCGTTCATTCTTCGAAAACGATCTGCGTTTCCTCAAACAGTTTAAA
TAAggaggacaatcatgaaaaagcctgaactcaccgcgacgtctgtcgagaagtttctgatcgaaaagttcgacagcgtctccgacctgatgcagctc
tcggagggcgaagaatctcgtgctttcagcttcgatgtaggagggcgtggatatgtcctgcgggtaaatagctgcgccgatggtttctacaaagatcgttat
gtttatcggcactttgcatcggccgcgctcccgattccggaagtgcttgacattggggaattcagcgagagcctgacctattgcatctcccgccgtgcaca
gggtgtcacgttgcaagacctgcctgaaaccgaactgcccgctgttctgcagccggtcgcggaggccatggatgcgatcgctgcggccgatcttagcc
agacgagcgggttcggcccattcggaccgcaaggaatcggtcaatacactacatggcgtgatttcatatgcgcgattgctgatccccatgtgtatcactgg
caaactgtgatggacgacaccgtcagtgcgtccgtcgcgcaggctctcgatgagctgatgctttgggccgaggactgccccgaagtccggcacctcgt
gcacgcggatttcggctccaacaatgtcctgacggacaatggccgcataacagcggtcattgactggagcgaggcgatgttcggggattcccaatacga
ggtcgccaacatcttcttctggaggccgtggttggcttgtatggagcagcagacgcgctacttcgagcggaggcatccggagcttgcaggatcgccgcg
gctccgggcgtatatgctccgcattggtcttgaccaactctatcagagcttggttgacggcaatttcgatgatgcagcttgggcgcagggtcgatgcgacg
caatcgtccgatccggagccgggactgtcgggcgtacacaaatcgcccgcagaagcgcggccgtctggaccgatggctgtgtagaagtacttgccgat
agtggaaaccgacgccccagcactcgtccgagggcaaaggaataacctaggGTTTAAACGAAGGCTCTCAAGGGCATCG
GTCGcacgcttgtg

Table S1. Plasmids used in this study.

| Plasmid Name | Description | Reference |
|---|---|---|
| Ara-pEffector-TS1 | CloDF13 ori smR J23119 crRNA-TS1 araC pBAD VchTniQ VchCas8 VchCas7 VchTnsA VchTnsB VchTnsC | This study |
| Ara-pEffector-TS2 | CloDF13 ori smR J23119 crRNA-TS2 araC pBAD VchTniQ VchCas8 VchCas7 VchTnsA VchTnsB VchTnsC | This study |
| Ara-pEffector-TS3 | CloDF13 ori smR J23119 crRNA-TS3 araC pBAD VchTniQ VchCas8 VchCas7 VchTnsA VchTnsB VchTnsC | This study |
| Ara-pEffector-TS4 | CloDF13 ori smR J23119 crRNA-TS4 araC pBAD VchTniQ VchCas8 VchCas7 VchTnsA VchTnsB VchTnsC | This study |
| Ara-pEffector-NT | CloDF13 ori smR J23119 crRNA-NT araC pBAD VchTniQ VchCas8 VchCas7 VchTnsA VchTnsB VchTnsC | Adapted from (1) |
| Ara-pEffector (no crRNA) | CloDF13 ori smR araC pBAD VchTniQ VchCas8 VchCas7 VchTnsA VchTnsB VchTnsC | Adapted from (1) |
| pTarget-TS1 | Heat sensitive pSC101 ori mScarlet PheS HygR J23119 crRNA-TS1 | This study |
| pTarget-TS2 | Heat sensitive pSC101 ori mScarlet PheS HygR J23119 crRNA-TS2 | This study |
| pTarget-TS3 | Heat sensitive pSC101 ori mScarlet PheS HygR J23119 crRNA-TS3 | This study |
| pTarget-TS4 | Heat sensitive pSC101 ori mScarlet PheS HygR J23119 crRNA-TS4 | This study |
| pKW20 | CloDF13 ori smR Lambda Red Alpha Gamma Exo Cas9 | Taken from (2) |
| **Location** | **Location** | **Reference** |
| TS1 | accagacccgcgagcattaattcttgcctcca | Designed from (2) |
| TS2 | tggtggcggtggtgggagctattctcgttctg | Designed from (2) |
| TS3 | gaaaacacctgatatgaaaggcaatgccacca | Designed from (2) |
| TS4 | agctgcaacaatgttgaaaatgccagccaact | Designed from (2) |
| NT | ctcctagcggtttgtgacaagtgtcagacaac | Designed from (1) |

Table S2. Primers used in this study.

| DNA oligonucleotides for linear transformations and genotyping | | |
|---|---|---|
| Oligo name | Sequence (5'-3') | Description |
| P-001 | CACGTGTTGACAATTAATCATCGGCAT | Universal primer for round one of nested PCR with EM7 promoter |
| P-002 | CCACAAAACAACCATATATTGATATCTCACAAAACAACCATAAGTTGATATTTTTGTGAATCGAGTATTTCAGCAAAACTACTGCAGTAAGatttgtcctactcaggagagcgttca | Round one of nested PCR to amplify EM7 3kb |
| P-003 | TGTTGATACAACCATAAAATGATAATTACACCCATAAATTGATAATTATCACACCCAcacgtgttgacaattaatcatcggcat | Universal primer for round two of nested PCR with EM7 promoter |
| P-004 | TGTTGATGCAACCATAAAGTGATATTTAATAATTATTTATAATCAGCAACTTAACCACAAAACAACCATATATTGATATCTCACAAAACAAC | Universal primer for round two of nested PCR |
| P-005 | CCACAAAACAACCATATATTGATATCTCACAAAACAACCATAAGTTGATATTTTTGTGAATCGAGTATTTCAGCAAAACTACTGCAGTAAGGCATTTTTTTCACTGCATTCTAGTTGTGG | Round one of nested PCR to amplify EM7 10kb |
| P-006 | ttgacagctagctcagtcctaggtataatactagtGGAGGACAATCATGGAGAAAAAAATCACTGG | Round one of nested PCR to amplify Jseries 1kb |
| P-007 | CCACAAAACAACCATATATTGATATCTCACAAAACAACCATAAGTTGATATTTTTGTGAATCGAGTATTTCAGCAAAACTACTGCAGTAAGttagctgtgttaCGCCCCGCC | Round one of nested PCR to amplify Jseries and EM7 1kb |
| P-008 | TGTTGATACAACCATAAAATGATAATTACACCCATAAATTGATAATTATCACACCCAttgacagctagctcagtcctaggt | Universal primer for round two of nested PCR with Jseries promoter |
| P-009 | ttgacagctagctcagtcctaggtataatactagtAAGGTGAGGAACTAAACCATGAACATCA | Round one of nested PCR for 3kb and 10kb product with Jseries promoter |
| P-010 | gggcgcggtagccgctgcgccctgtcaatttcccttccttattagccgctCACGTGTTGACAATTAATCATCGGCAT | Lambda Red 3kb and 10kb product for TS1 integration (EM7 driven) |
| P-011 | gatatcagcgaaaacataagcaaaagtgaatgttttaagaacattccgtaGCATTTTTTTCACTGCATTCTAGTTGTGG | Lambda Red 10kb product for TS1 integration |
| P-012 | gcatgaagtatggcaaaaaggggctgatgaaaatgccagcacaatacgatCACGTGTTGACAATTAATCATCGGCAT | Lambda Red 3kb and 10kb product for TS4 integration (EM7 driven) |
| P-013 | aatgagtaaaatcttatgtttaatttatttgtggggttttaattaatttaGCATTTTTTTCACTGCATTCTAGTTGTGG | Lambda Red 10kb product for TS4 integration |
| P-014 | gatatcagcgaaaacataagcaaaagtgaatgttttaagaacattccgtaATTTGTCCTACTCAGGAGAGCGTTCA | Lambda Red 3kb product for TS1 integration |
| P-015 | aatgagtaaaatcttatgtttaatttatttgtggggttttaattaatttaATTTGTCCTACTCAGGAGAGCGTTCA | Lambda Red 3kb product for TS4 integration |
| P-016 | GGGCGCGGTAGCCGCTGCGCCCTGTCAATTTCCCTTCCTTATTAGCCGCTttgacagctagctcagtcctaggtataatactagtGGAGGACAATCATGGAGAAAAAAATCACTGG | Lambda Red 1kb product for TS1 integration (Jseries driven) |
| P-017 | GCATGAAGTATGGCAAAAAGGGGCTGATGAAAATGCCAGCACAATACGATttgacagctagctcagtcctaggtataatactagtGGAGGACAATCATGGAGAAAAAAATCACTGG | Lambda Red 1kb product for TS4 integration (Jseries driven) |
| P-018 | gggcgcggtagccgctgcgccctgtcaatttcccttccttattagccgctTTGACAGCTAGCTCAGTCCTAGGTATAATACTAGTaaggtgaggaactaaaccATGAACATCAA | Lambda Red 3kb and 10kb product for TS1 integration (Jseries driven) |
| P-019 | gcatgaagtatggcaaaaaggggctgatgaaaatgccagcacaatacgatTTGACAGCTAGCTCAGTCCTAGGTATAATACTAGTaaggtgaggaactaaaccATGAACATCAA | Lambda Red 3kb and 10kb product for TS4 integration (Jseries driven) |

| | | |
|---|---|---|
| P-020 | gagttcaagcgtaagaacgaaagccaactg | TS1 genome integration check |
| P-021 | gcgatgagtgatgaataacgaccattacagc | TS1 genome integration check |
| P-022 | acctgtattctcgggatagtcggtg | TS2 genome integration check |
| P-023 | ttcacccgtaaagcaggtgtcgagta | TS2 genome integration check |
| P-024 | aatgatgctcgattttctcggcaatgg | TS3 genome integration check |
| P-025 | gaaacggaaacaataaaaggtgaaatgacgc | TS3 genome integration check |
| P-026 | gattcacaggcggatcgcggtctata | TS4 genome integration check |
| P-027 | tcatgcgaccgtcatcaatgtgag | TS4 genome integration check |
| **DNA oligonucleotides for Plasmid Construction** | | |
| Oligo name | Sequence (5'-3') | Description |
| P-028 | GGATCCGAATTCGAGCGAAGGAGATATACATATG | Construction of Ara-pEffector (backbone) |
| P-029 | actagtattatacctaggactgagctagctgtc | Construction of Ara-pEffector (backbone) |
| P-030 | tcgagagatccggctgctaaca | Construction of Ara-pEffector (araC and pBad insert) |
| P-031 | ATCTCCTTCGCTCGAATTCGGATCCcgggtatggagaaacagtagagagttgcga | Construction of Ara-pEffector (araC and pBad insert) |
| P-032 | ctcagtcctaggtataatactagtCAcGTGAACTGCCGAGTAGGTAGCTGATAACaccagacccgcgagcattaattcttgcctcca | Construction of Ara-pEffector (TS1 crRNA) |
| P-033 | gtttgttagcagccggatctctcgaGTTATCAGCTACCTACTCGGCAGTTCACtggaggcaagaattaatgctcgcgggtctggt | Construction of Ara-pEffector (TS1 crRNA) |
| P-034 | ctcagtcctaggtataatactagtCAcGTGAACTGCCGAGTAGGTAGCTGATAACtggtggcggtggtgggagctattctcgttctg | Construction of Ara-pEffector (TS2 crRNA) |
| P-035 | gtttgttagcagccggatctctcgaGTTATCAGCTACCTACTCGGCAGTTCACcagaacgagaatagctcccaccaccgccacca | Construction of Ara-pEffector (TS2 crRNA) |
| P-036 | ctcagtcctaggtataatactagtCAcGTGAACTGCCGAGTAGGTAGCTGATAACgaaaacacctgatatgaaaggcaatgccacca | Construction of Ara-pEffector (TS3 crRNA) |
| P-037 | gtttgttagcagccggatctctcgaGTTATCAGCTACCTACTCGGCAGTTCACtggtggcattgcctttcatatcaggtgttttc | Construction of Ara-pEffector (TS3 crRNA) |
| P-038 | ctcagtcctaggtataatactagtCAcGTGAACTGCCGAGTAGGTAGCTGATAACagctgcaacaatgttgaaaatgccagccaact | Construction of Ara-pEffector (TS4 crRNA) |
| P-039 | gtttgttagcagccggatctctcgaGTTATCAGCTACCTACTCGGCAGTTCACagttggctggcattttcaacattgttgcagct | Construction of Ara-pEffector (TS4 crRNA) |
| P-040 | ctcagtcctaggtataatactagtCAcGTGAACTGCCGAGTAGGTAGCTGATAACctcctagcggtttgtgacaagtgtcagacaac | Construction of Ara-pEffector (NT crRNA) |
| P-041 | gtttgttagcagccggatctctcgaGTTATCAGCTACCTACTCGGCAGTTCACgttgtctgacacttgtcacaaaccgctaggag | Construction of Ara-pEffector (NT crRNA) |
| P-042 | CGACCGATGCCCTTGAGAGCCTT | Construction of Ara-pEffector (-crRNA) (backbone) |
| P-043 | TGAAGGCTCTCAAGGGCATCGGTCGcacgcttgtgaaacccatctaattg | Construction of Ara-pEffector (-crRNA) (backbone) |
| P-044 | GGATCCGAATTCGAGCGAAGGAGATATACATAT | Construction of Ara-pEffector (-crRNA) (araC and pBad insert) |
| P-045 | atctccttcgctcgaattcggatccCGGGTATGGAGAAACAGTAGAGAGTTGC | Construction of Ara-pEffector (-crRNA) (araC and pBad insert) |
| P-046 | ctccacaaaaggcgtagtgtacagtgaGTTAACcgaggcagggggaattcccaattagatg | Universal primer to insert crRNA into pTarget |
| P-047 | gtccgagggcaaaggaataacctaggGTTTAAACGAAGGCTCTCAAGGGCATCGGTCG | Universal primer to insert crRNA into pTarget |
| P-048 | GTTTAAACcctaggttattcctttgccct | pTarget backbone |
| P-049 | GTTAACtcactgtacactacgccttttg | pTarget backbone |

Table S3. Genomic target sites taken from (2) and equivalent names in this study.

| Location from (2) | Name in this Study |
|---|---|
| L02 | TS1 |
| L09 | TS2 |
| L14 | TS3 |
| L21 | TS4 |
